# Risk factors, prognostic factors, and nomograms for distant metastasis in patients with triple-negative breast cancer: a population-based study

**DOI:** 10.1101/2024.11.08.622627

**Authors:** Hongguo Guo, Song Qiao, Cai Cheng, Jun Liu, Shangzhen Yang, Wanling Lu

**Affiliations:** Department of Oncology, The 986th Hospital of the People’s Liberation Army Air Force,Xi’an,China

**Author notes:** Corresponding author (LWL). All authors contributed equally to this work.

**Keywords:** Triple-negative breast cancer, distant metastasis, nomograms, predictor, prognosis

## Abstract

**Background:** Triple-negative breast cancer (TNBC) is associated with a poor prognosis, higher invasiveness compared to other breast cancer variants, and limited treatment options. Due to the lack of conventional treatment methods, TNBC patients are prone to distant metastasis. The purpose of this study is to evaluate the clinical and pathological characteristics of TNBC patients, predict the impact of risk factors on their prognosis, and establish a predictive model to assist doctors in treating TNBC patients and improving their outcomes.

**Methods:** We conducted a retrospective study on patients with TNBC from 2010 to 2015 in the Surveillance, Epidemiology, and End Results (SEER) database. We gathered tumor characteristics and treatment data from these patients. Univariate and multivariate logistic regression analyses were employed to identify independent risk factors for distant metastasis in TNBC patients, while univariate and multivariate Cox proportional hazards regression analyses were utilized to determine independent prognostic factors for TNBC patients with distant metastasis. Based on these factors, a new nomogram was constructed, and patients were categorized into high and low-risk groups using the nomogram scores. The prognosis of the patients was analyzed using Kaplan-Meier (K-M) survival analysis.

**Results:** This study enrolled a total of 16,959 patients with TNBC, among whom 730 (4.3%) had distant metastasis at the time of diagnosis. Independent risk factors for distant metastasis in TNBC patients encompass being married, having an invasive lobular carcinoma (ILC) pathology, advanced T stage, advanced N stage, no radiotherapy, no surgery, and tumor size exceeding 50mm. The area under the curve (AUCs) for the training set and validation set were 0.892 and 0.907, respectively. The independent prognostic factors for patients with distant metastasis were being unmarried (P < 0.001), not undergoing surgery (P < 0.001), and not receiving chemotherapy (P < 0.001). The AUCs for the training set at 12, 36, and 60 months were 0.751, 0.709, and 0.691, respectively, while the AUCs for the validation set at 12, 36, and 60 months were 0.690, 0.690, and 0.732, respectively. The AUCs for the extended test set at 12, 36, and 60 months were 0.741, 0.727, and 0.718 respectively. The results of the calibration curve, decision curve analysis (DCA), and K-M survival curve confirm that both nomograms can accurately predict the occurrence of distant metastasis and prognosis in TNBC patients.

**Conclusion:** We constructed two nomograms to predict distant metastasis risk in TNBC patients and prognosis in patients with distant metastasis. Our findings suggest that they could serve as effective predictive tools and may assist in clinical decision-making.

## Introduction

In the realm of female cancers, breast cancer boasts the highest mortality rate, with an estimated 2.3 million new cases (11.7%) and 685,000 deaths (6.9%) in 2020 [1]. Triple-Negative Breast Cancer (TNBC) stands as a distinct subtype of breast cancer, marked by the absence of expression for estrogen receptor (ER), progesterone receptor (PR), and human epidermal growth factor receptor 2 (HER2). Statistical data indicates that TNBC comprises 10% to 20% of all breast cancer cases[2]. In comparison to other forms of breast cancer, TNBC exhibits more intricate biological traits, frequently manifesting as higher tumor malignancy and a poorer prognosis.

Because ER, PR and HER 2 are not expressed, the prognosis is poor, the treatment options are less, and the traditional targeted therapy and endocrine therapy are ineffective[3]. Subjects with TNBC usually only rely on conventional chemotherapy methods, and this treatment regimen may not be effective in controlling the tumor progression. Studies show that unlike other breast cancer variants, TNBC has higher 5-year mortality, is more aggressive, has a poorer prognosis, and reaches peak recurrence in the first 3 years of diagnosis. Distant metastasis occurs in about 50% of patients[4,5], and survival is even lower for patients with locally advanced or metastatic TNBC.

TNBC patients are more prone to distant metastasis, and early detection of distant metastasis through screening can improve survival rates.[6]. For a long time, the number of studies on the relationship between traditional clinical pathological features and distant metastasis in TNBC has been limited. More research has focused on metastatic tumors, while overlooking a comprehensive understanding of non-metastatic cases.[7]. Moreover, the prognosis of patients who have already experienced distant metastasis at the time of diagnosis requires further study. Risk factors associated with TNBC distant metastasis may identify high-risk patients, thereby aiding early clinical intervention.

Many years have passed since the American Joint Committee on Cancer (AJCC) prognostic staging system for tumors, lymph nodes, and metastasis (TNM) was first introduced to determine the prognosis of breast cancer patients. With the development of molecular typing and precision therapies, the importance of traditional tumor staging methods gradually decreases. Therefore, based on these data does not fully evaluate the possibility and future prospects of distant metastasis in TNBC patients[8,9] Due to the convenience and accuracy of nomograms, it has recently been widely generated to evaluate the prognosis of cancer patients, which is a good choice [10,11]. Therefore, this study recruited a large number of TNBC cases for analysis using the Surveillance, Epidemiology, and End Results (SEER) database. We investigated demographic characteristics (age, race, and marital status), tumor characteristics (histological type, grade, clinical T stage and N stage, tumor size) and treatment (surgery, chemotherapy, radiotherapy) to assess the incidence, risk factors, and prognosis of TNBC metastasis, And developed two nomograms, which help clinicians make more appropriate medical decisions.

## Materials and Methods

### Source of the data and patient selection

The study used a database called SEER, which was released in March 2024. Patients included in the study were selected from SEER * Stat version 8.4.3, the database contains population, patient characteristics, tumor characteristics, diagnosis and healthcare information from 17 cancer registries, representing approximately 28% of cancer cases in the United States between 2000 and 2021. A signed SEER study data agreement form was submitted to the SEER project for access and analysis of the SEER database (https://seer.cancer.gov/). Given that reporting of cancer reports is mandatory in every state across the country, obtaining patient informed consent is not important to access the SEER database.

Based on the following inclusion criteria, an analysis was conducted on breast cancer patients from January 1, 2010 to December 31, 2015 (1): female 18 to 90 years old; (2) initial diagnosis of primary breast cancer and (3) pathological analysis (ER- / PR- / HER 2-) confirming the triple-negative molecular subtype.(4) demographic variables, including age, sex and race are available; (5) clinicopathological information, including primary tumor site, grade, histological type, TNM and tumor size are available. Exclusion criteria included: (1) patients with multiple primary tumors, and (2) patients with missing or incomplete follow-up information. Data entered into the case table included age at diagnosis, ethnicity, marital status, histological type, grade, AJCC stage, tumor size, cancer treatment, etc. Relevant data factors were analyzed, and participants lacking clinical characteristics or survival information were excluded from the final dataset, and finally this study included 16,959 TNBC patients, including 730 patients with distant metastases.

The age classification in this paper follows the 2023 United Nations World Health Organization guidelines, which are based on the assessment of global human body mass and average life expectancy. Accordingly, those aged 18-44 were considered young; those aged 45 to 59 were classified as middle-aged; and those aged over 60 were classified as elderly. Many studies have established that morphologic evaluation of differentiation can provide valuable prognostic insights in breast cancer. Specifically, grade I tumors were well differentiated, grade II moderately differentiated, grade III poorly differentiated, and grade IV undifferentiated. It has been documented that for some anatomical sites, grades III and IV can be combined into a single level and that this classification is applicable for breast cancer. All patients were used to form a diagnostic cohort to explore risk factors for distant metastases and establish a predictive nomogram. Furthermore, available specific treatment information, including surgery, chemotherapy and radiotherapy, form prognostic cohorts to study prognostic factors in distant metastases patients and develop new prognostic nomograms.

Patients were divided into diagnostic and prognostic cohorts. In the diagnostic cohort, patients were randomly assigned to a training set (70%) and a validation set (30%), with a ratio of 7:3. In the prognostic cohort, it was composed of patients with distant metastasis from both the training and validation sets of the diagnostic cohort. For each cohort, to prevent overfitting and ensure accuracy, nomograms were constructed using patients from the training set and validated using patients from the validation set.

### Data collection

In this study, age, race, marital status, pathological type, grade, T stage, N stage, primary site surgery, radiotherapy, chemotherapy, and tumor size were used as variables for risk factors of distant metastasis in TNBC patients. Additionally, we conducted survival analyses with overall survival (OS) as the primary outcome to explore prognostic factors for patients with distant metastasis.

### Statistical analysis

All studies were analyzed using R software (version 4.3.3), and a P<0.05 (two-sided) considered statistical significance. All patients were randomly divided into training and validation sets, and the distribution of two group variables was compared by chi-square test or Fishers exact test.

In the diagnostic cohort, univariate logistic analysis was employed to identify risk factors associated with distant metastasis. Variables with P<0.05 in the univariate analysis were subjected to multivariate logistic analysis to determine independent risk factors for distant metastasis in patients[12]. Additionally, nomograms and receiver operating characteristic (ROC) curves were generated for all independent variables based on individual risk factors, and the corresponding area under the curve (AUC) was calculated to assess their discriminative ability. Additionally, calibration curves and decision curve analysis (DCA) were used to evaluate the performance of the nomograms.

In terms of prognostic factors, we used univariate Cox regression analysis to identify factors associated with overall survival (OS) in patients with distant metastases. Subsequently, significant variables with P<0.05 were subjected to "stepwise" multivariate Cox analysis to further determine independent prognostic factors. We constructed a prognostic nomogram based on these independent prognostic predictors to predict the overall survival of patients with distant metastases and calculated individual risk scores using the nomogram formula. Additionally, the concordance index (C-index) was used to assess the probability of agreement between actual outcomes and predicted outcomes. Time-dependent receiver operating characteristic (ROC) curves were generated for the nomogram and all independent prognostic variables at 12, 36, and 60 months, and the time-dependent area under the curve (AUC) was applied to evaluate the discriminative ability of the prognostic model. Calibration curves and decision curve analysis (DCA) were plotted at 12, 36, and 60 months to assess the flexibility and clinical applicability of the predictive model. All patients with distant metastases were divided into high-risk and low-risk groups based on the median risk score. Kaplan-Meier (K-M) survival curves and log-rank tests were used to demonstrate the difference in overall survival status between the two groups.

The aforementioned statistical analysis utilized various R packages such as "Survival", "forestplot", "Glmnet", "rms", "stdca.R", "survivalROC", and "survivalminer" (http://www.r-project.org/).

## Results

### Baseline characteristics of the study population

A total of 16,959 TNBC patients who met the criteria were included from the SEER database, of which 11,871 patients were included in the training set and 5,088 patients in the validation set. The average ages for the training and validation sets were 55.95 years (ranging from 20 to 90 years) and 56.16 years (ranging from 21 to 90 years), respectively. The most common pathological type was invasive ductal carcinoma (IDC) (86.3% for the training set and 86.9% for the validation set), and the most common tumor differentiation grade was III – IV (81.1% for the training set and 81.3% for the validation set). The most common T and N stages were T1 (44.4% for the training set and 45.0% for the validation set) and N0 (65.4% for the training set and 65.1% for the validation set), respectively. Overall, no significant differences were found between the training and validation sets (P > 0.05). (Table 1).

**Table 1.**
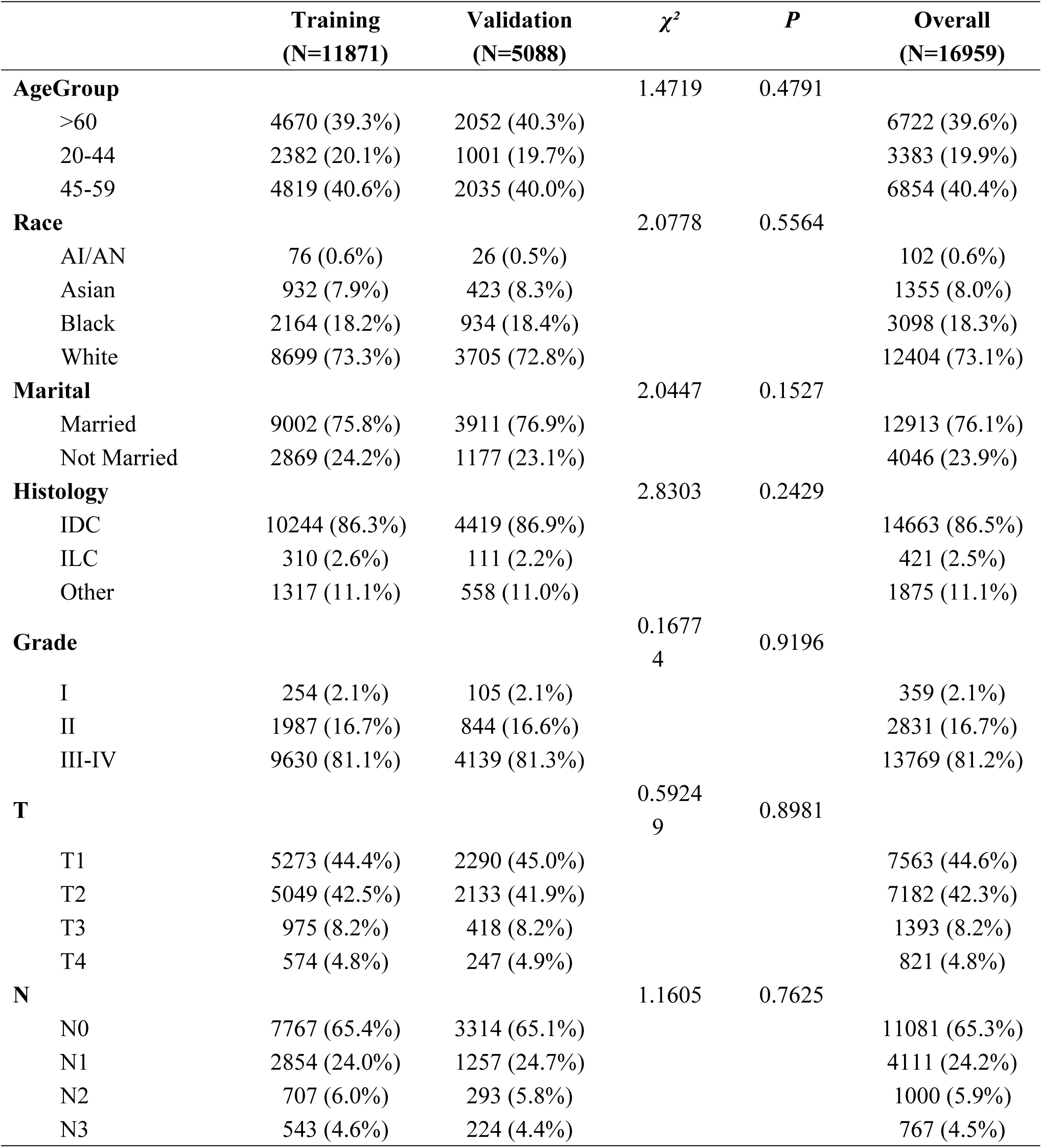
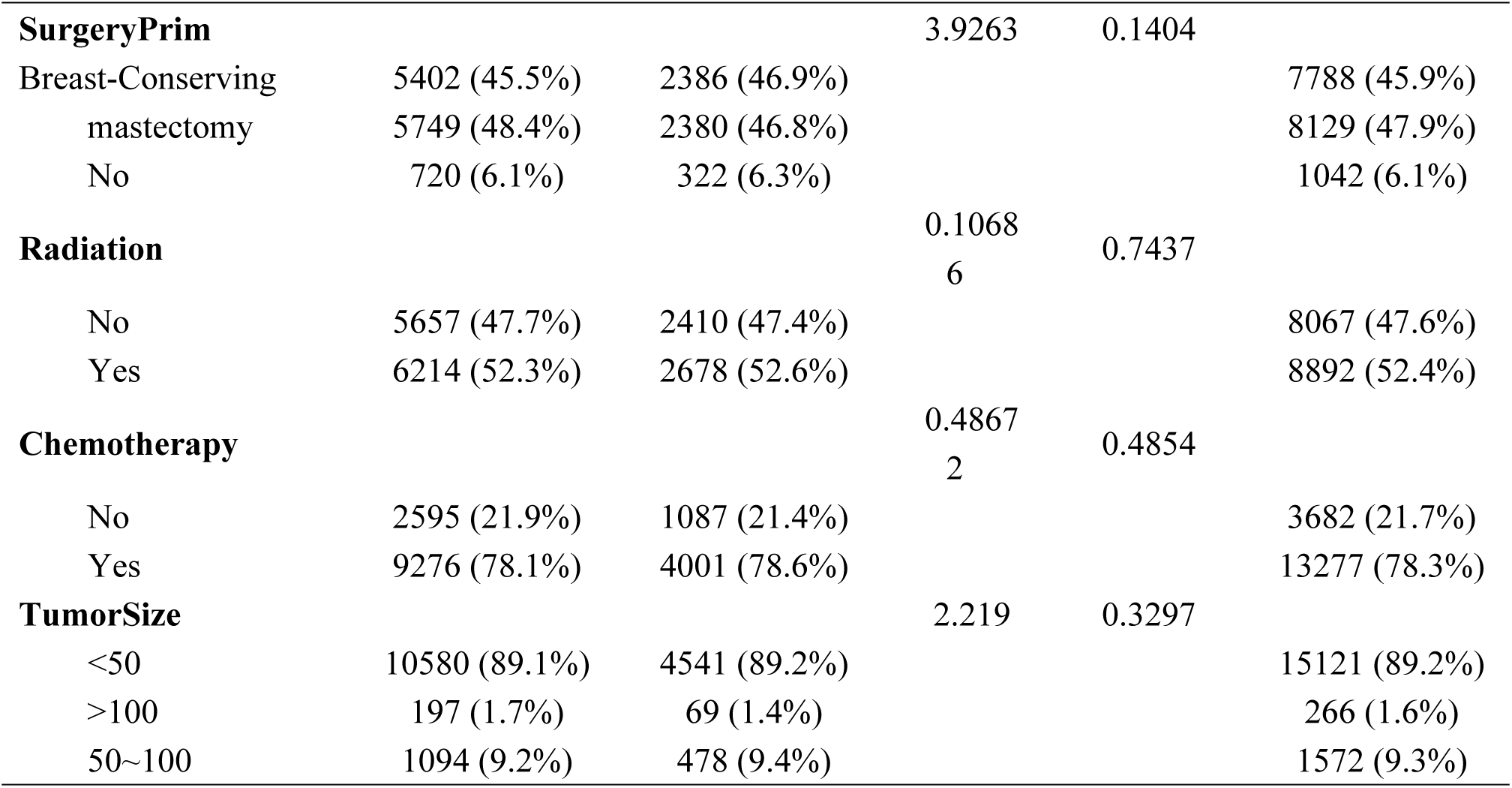
Baseline clinical characteristics of TNBC patients.

### Incidence and risk factors for distant metastasis in patients with TNBC

At the initial diagnosis, 730 cases (4.3%) were identified with distant metastasis, whereas 16,229 cases (95.7%) were found to be free of it. Table 2 presents the results of a univariate logistic regression analysis conducted on 11 potential factors, revealing that eight variables are significantly associated with distant metastasis. These variables include marital status, pathological type, T stage, N stage, primary site surgery, radiotherapy, chemotherapy, and tumor size. Furthermore, a multivariate logistic regression analysis has pinpointed several independent risk predictors for distant metastasis in TNBC patients: being married,pathological invasive lobular carcinoma (ILC), high T stage,high N stage, no radiotherapy, no surgery, and tumor size exceeding 50 mm (Table 2).

**Table 2.** Univariate and multivariate logistic analysis of distant metastasis in TNBC patients.

|  | Univariate analysis |  |  | Multivariate analysis |  |  |
| --- | --- | --- | --- | --- | --- | --- |
|  | OR | 95%CI | P | OR | 95%CI | P |
| **AgeGroup** |  |  |  |  |  |  |
| 20-44 | Reference |  |  |  |  |  |
| 45-59 | 0.965 | 0.814-1.148 | 0.735 |  |  |  |
| >60 | 1.065 | 0.900-1.265 | 0.540 |  |  |  |
| **Race** |  |  |  |  |  |  |
| White | Reference |  |  |  |  |  |
| Black | 1.165 | 0.994-1.359 | 0.107 |  |  |  |
| Asian | 0.932 | 0.726-1.179 | 0.633 |  |  |  |
| AI/AN | 1.949 | 0.999-3.428 | 0.072 |  |  |  |
| **Marital** |  |  |  |  |  |  |
| Married | Reference |  |  | Reference |  |  |
| Not Married | 1.457 | 1.271-1.666 | <0.001 | 0.814 | 0.690-0.958 | 0.039 |
| **Histology** |  |  |  |  |  |  |
| IDC | Reference |  |  | Reference |  |  |
| ILC | 2.350 | 1.743-3.105 | <0.001 | 1.843 | 1.298-2.571 | 0.003 |
| Other | 1.306 | 1.083-1.565 | 0.017 | 1.017 | 0.807-1.272 | 0.900 |
| **Grade** |  |  |  |  |  |  |
| I | Reference |  |  |  |  |  |
| II | 1.617 | 0.945-3.028 | 0.171 |  |  |  |
| III-IV | 1.797 | 1.073-3.315 | 0.085 |  |  |  |
| **T** |  |  |  |  |  |  |
| T1 | Reference |  |  | Reference |  |  |
| T2 | 2.576 | 2.121-3.145 | <0.001 | 1.554 | 1.262-1.923 | <0.001 |
| T3 | 8.783 | 7.069-10.941 | <0.001 | 1.774 | 1.191-2.632 | 0.017 |
| T4 | 30.340 | 24.706-37.435 | <0.001 | 5.044 | 3.671-6.916 | <0.001 |
| **N** |  |  |  |  |  |  |
| N0 | Reference |  |  | Reference |  |  |
| N1 | 5.873 | 4.995-6.925 | <0.001 | 3.021 | 2.511-3.638 | <0.001 |
| N2 | 6.714 | 5.342-8.403 | <0.001 | 4.252 | 3.261-5.520 | <0.001 |
| N3 | 19.071 | 15.680-23.205 | <0.001 | 8.012 | 6.321-10.155 | <0.001 |
| **SurgeryPrim** |  |  |  |  |  |  |
| Breast-Conserving | Reference |  |  | Reference |  |  |
| mastectomy | 2.776 | 2.288-3.388 | <0.001 | 1.142 | 0.924-1.420 | 0.305 |
| No | 41.499 | 34.098-50.820 | <0.001 | 13.636 | 10.824-17.259 | <0.001 |
| **Radiation** |  |  |  |  |  |  |
| No | Reference |  |  | Reference |  |  |
| Yes | 0.493 | 0.433-0.561 | <0.001 | 0.633 | 0.537-0.746 | <0.001 |
| **Chemotherapy** |  |  |  |  |  |  |
| No | Reference |  |  |  |  |  |
| Yes | 1.400 | 1.189-1.658 | 0.001 |  |  |  |
| **TumorSize** |  |  |  |  |  |  |
| <50 | Reference |  |  | Reference |  |  |
| >100 | 13.865 | 10.898-17.533 | <0.001 | 2.343 | 1.607-3.417 | <0.001 |
| 50~100 | 5.941 | 5.147-6.847 | <0.001 | 1.803 | 1.305-2.503 | 0.003 |

### Nomogram and validation of TNBC diagnostic cohort

Based on these seven independent predictive factors, a nomogram was developed to predict the risk of distant metastasis in patients with TNBC (Fig 1). To validate the performance of the nomogram, ROC curves were plotted for both the training and validation sets, with the AUC being 0.892 and 0.907, respectively (Figs 2 and 3). To verify its predictive ability, ROC curves for all independent predictive factors were plotted (Figs 4 and 5), showing that their discriminative ability in the training and validation sets was superior to other single factors. At the same time, the calibration curves drawn showed good consistency between observed outcomes and predicted outcomes (Figs 6 and 7). By examining the DCA curves, the net benefit of using the scale as a tool to trigger medical intervention compared to comprehensive treatment or no treatment was demonstrated, allowing us to decide whether using the model to assist in clinical decision-making would improve the treatment outcomes of our patients (Figs 8 and 9).

**Fig 1.**
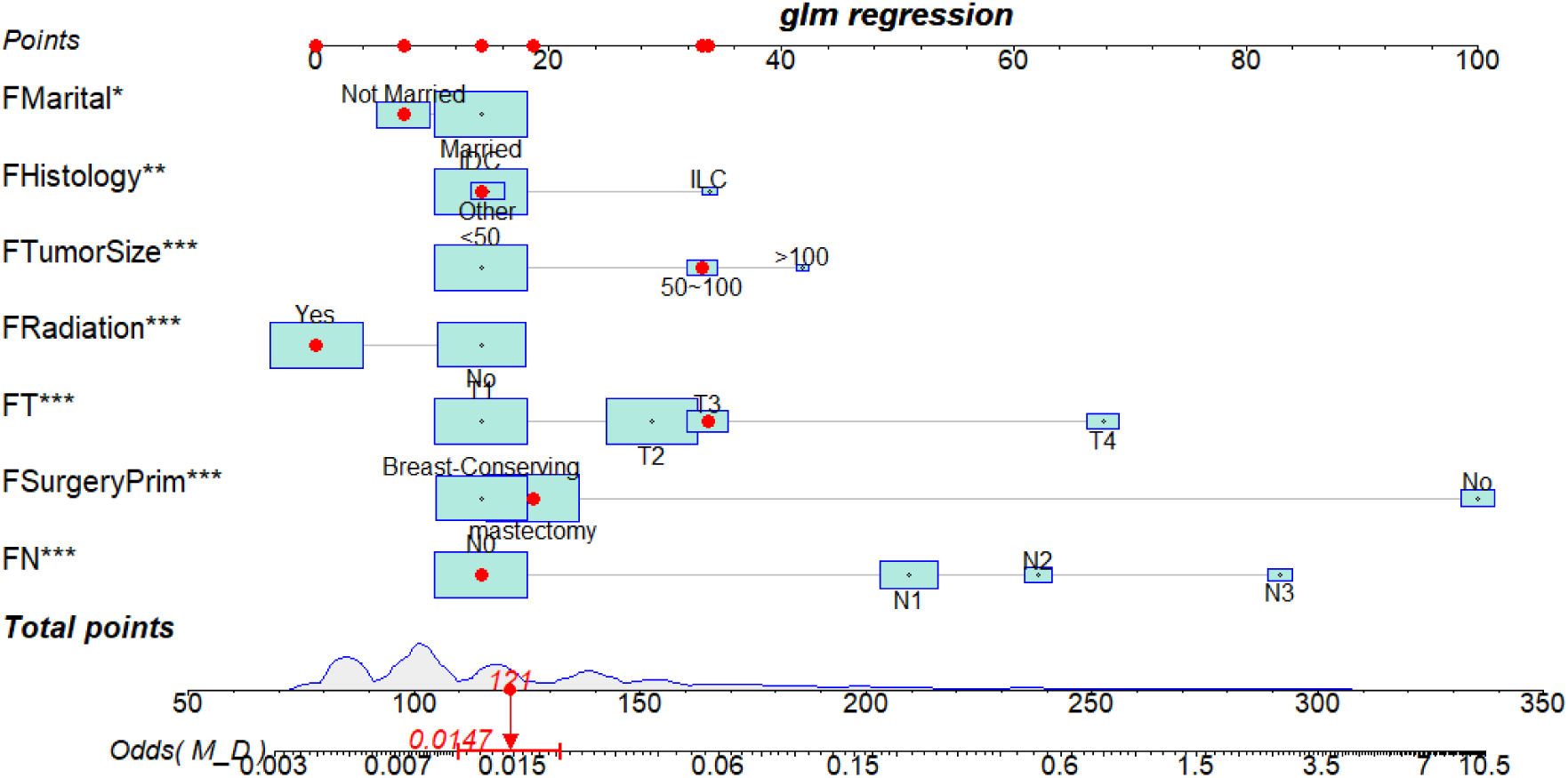
Nomogram for assessing the risk of distant metastasis in TNBC patients

**Fig 2.**
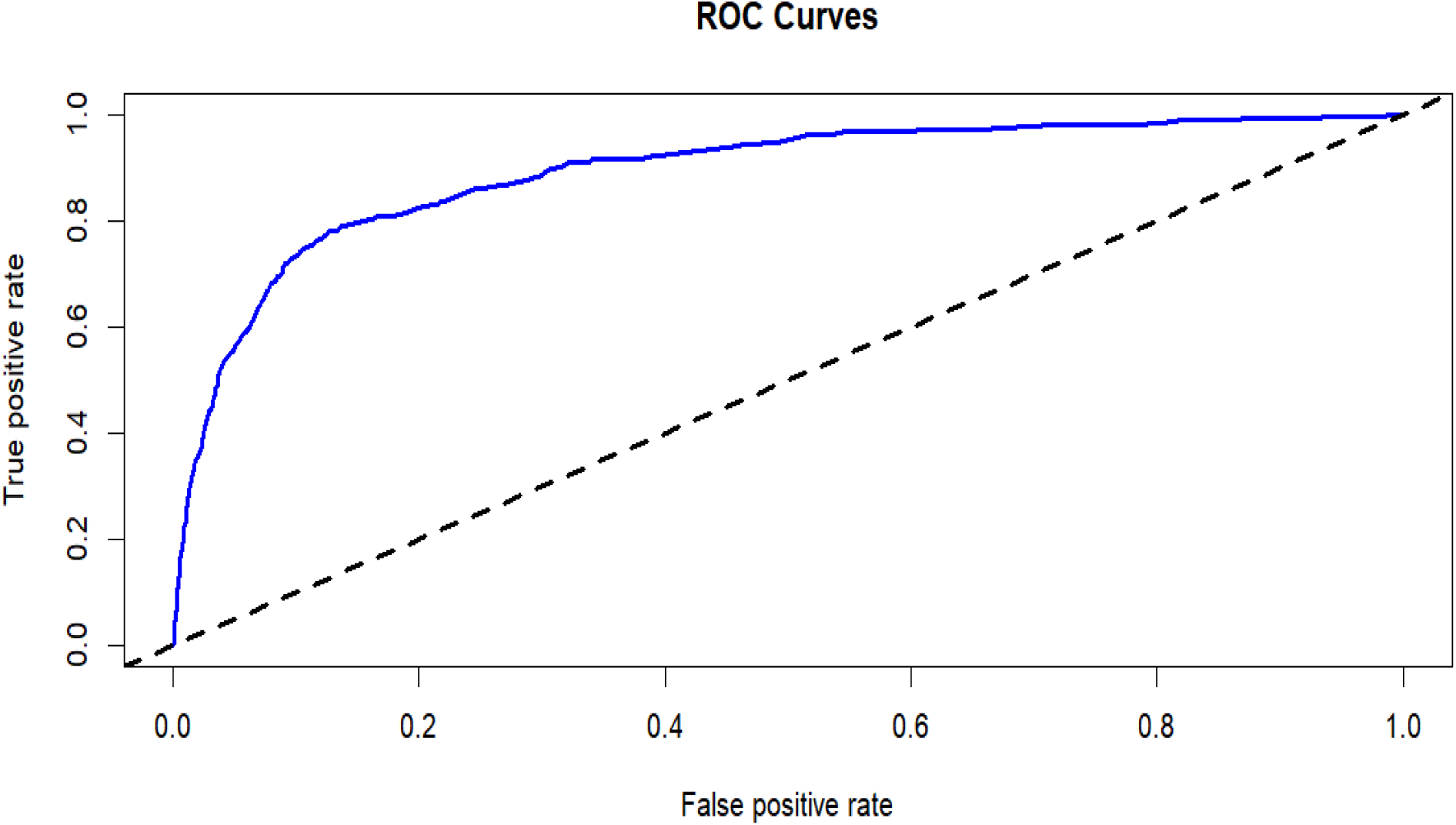
Receiver Operating Characteristic Curve for the Training Set AUC= 0.892

**Fig 3.**
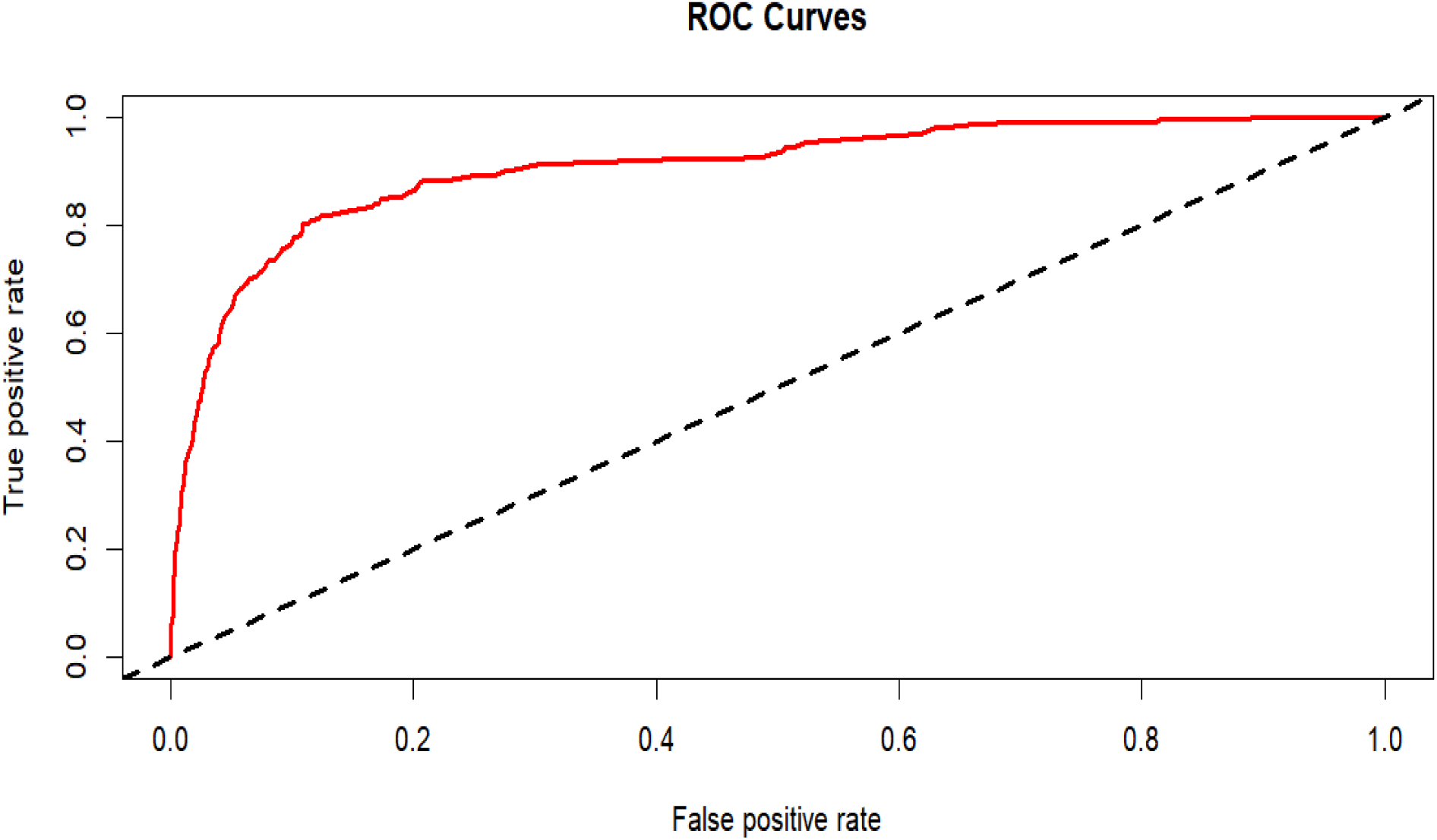
Receiver Operating Characteristic Curve for the Validation Set AUC= 0.907

**Fig 4.**
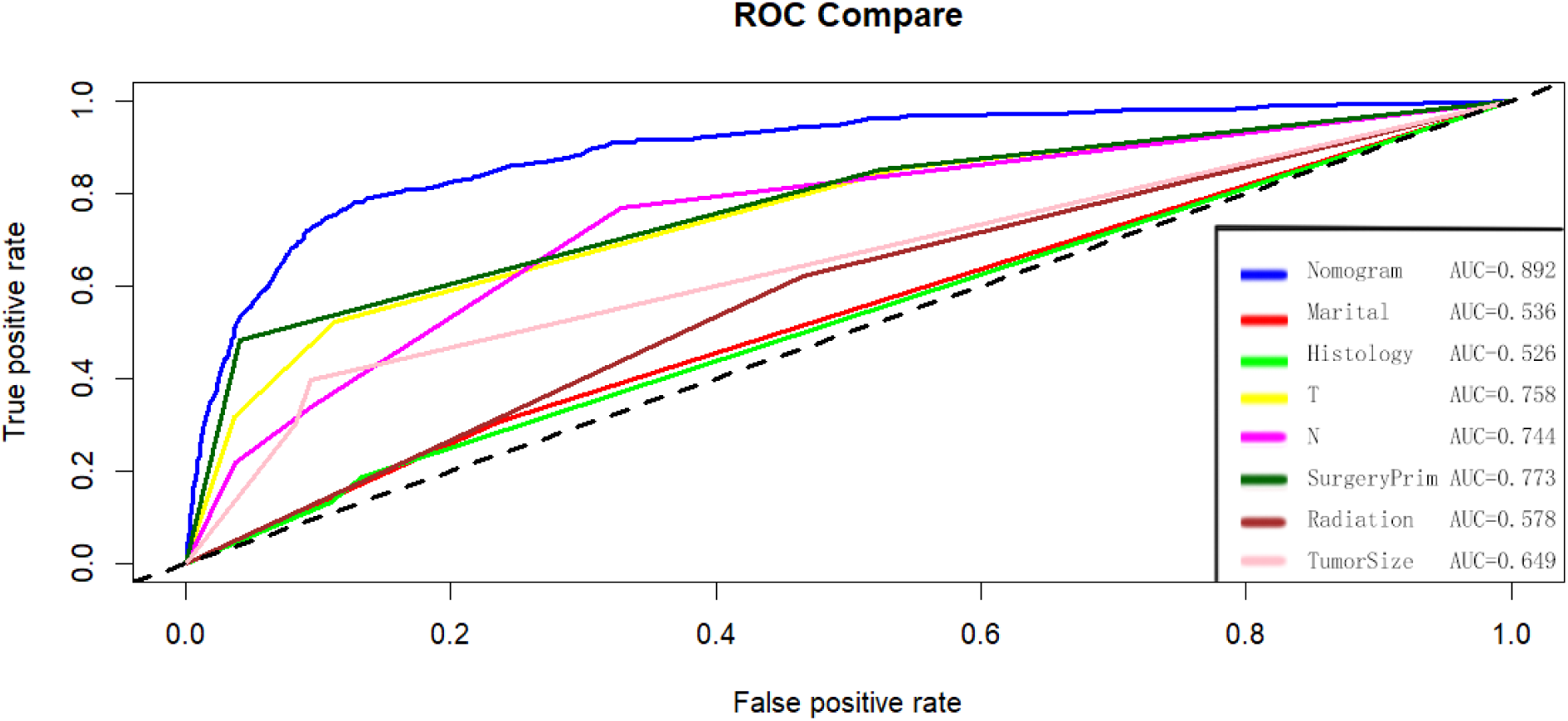
Comparison of the areas under the receiver operating characteristic curves for various factors in the training set

**Fig 5.**
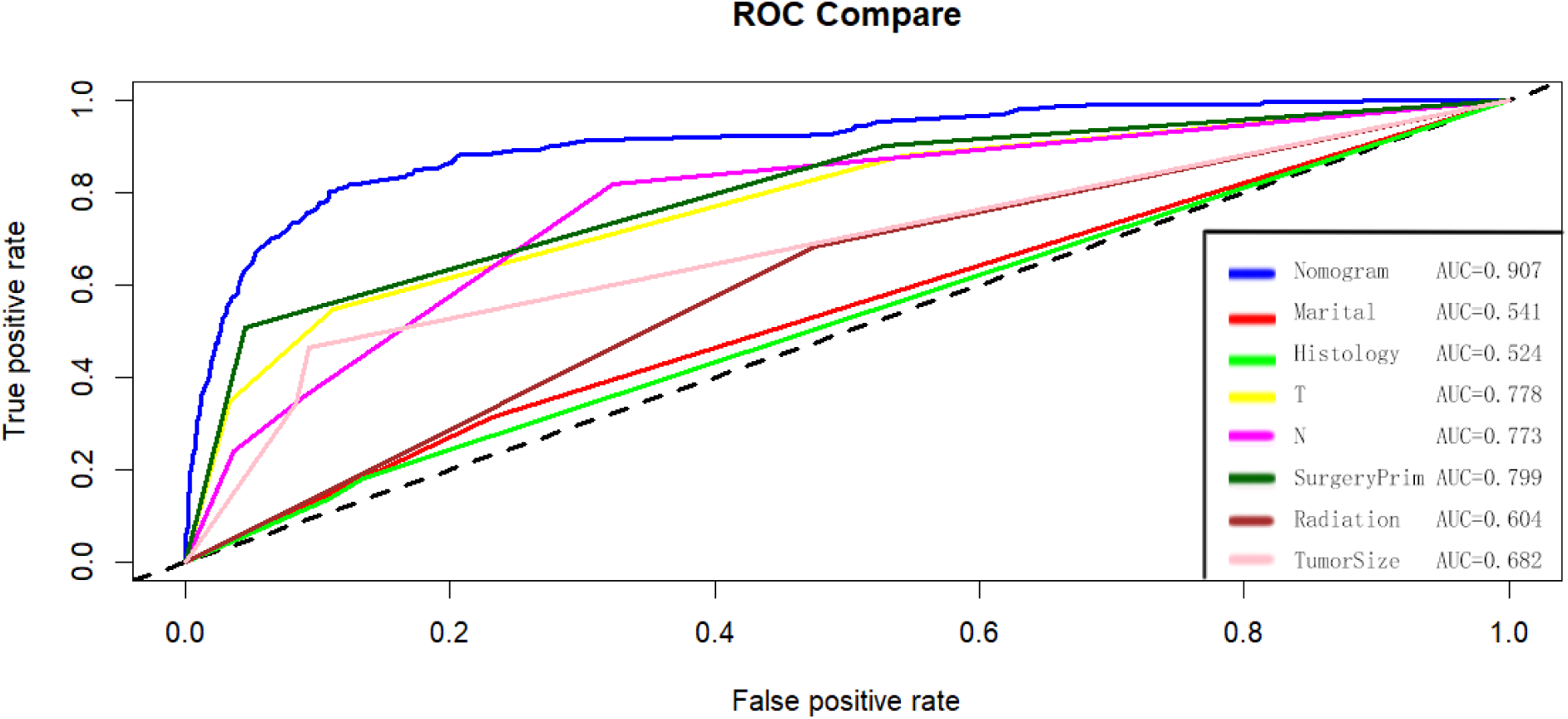
Comparison of the areas under the receiver operating characteristic curves for various factors in the Validation set

**Fig 6.**
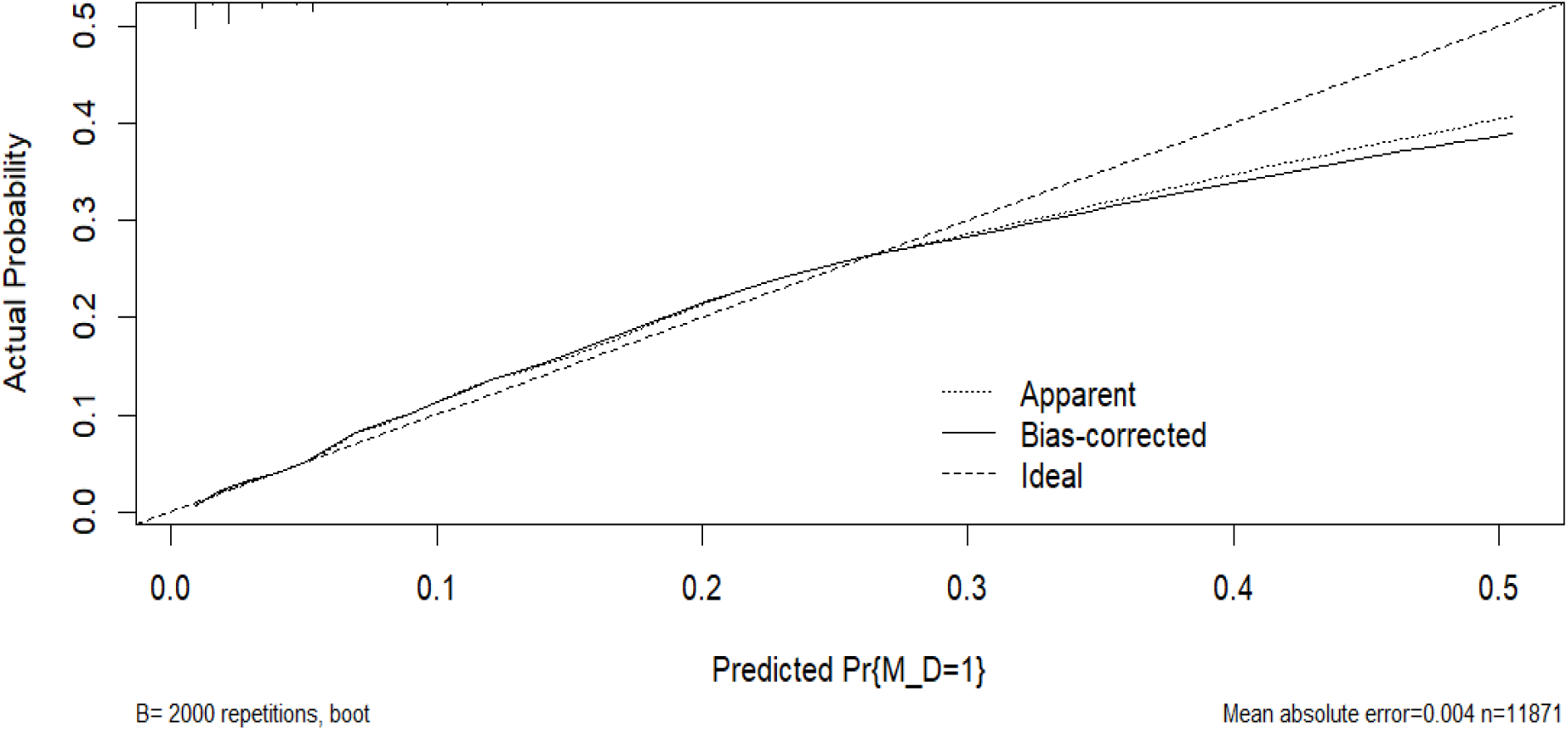
Calibration curve of the training set

**Fig 7.**
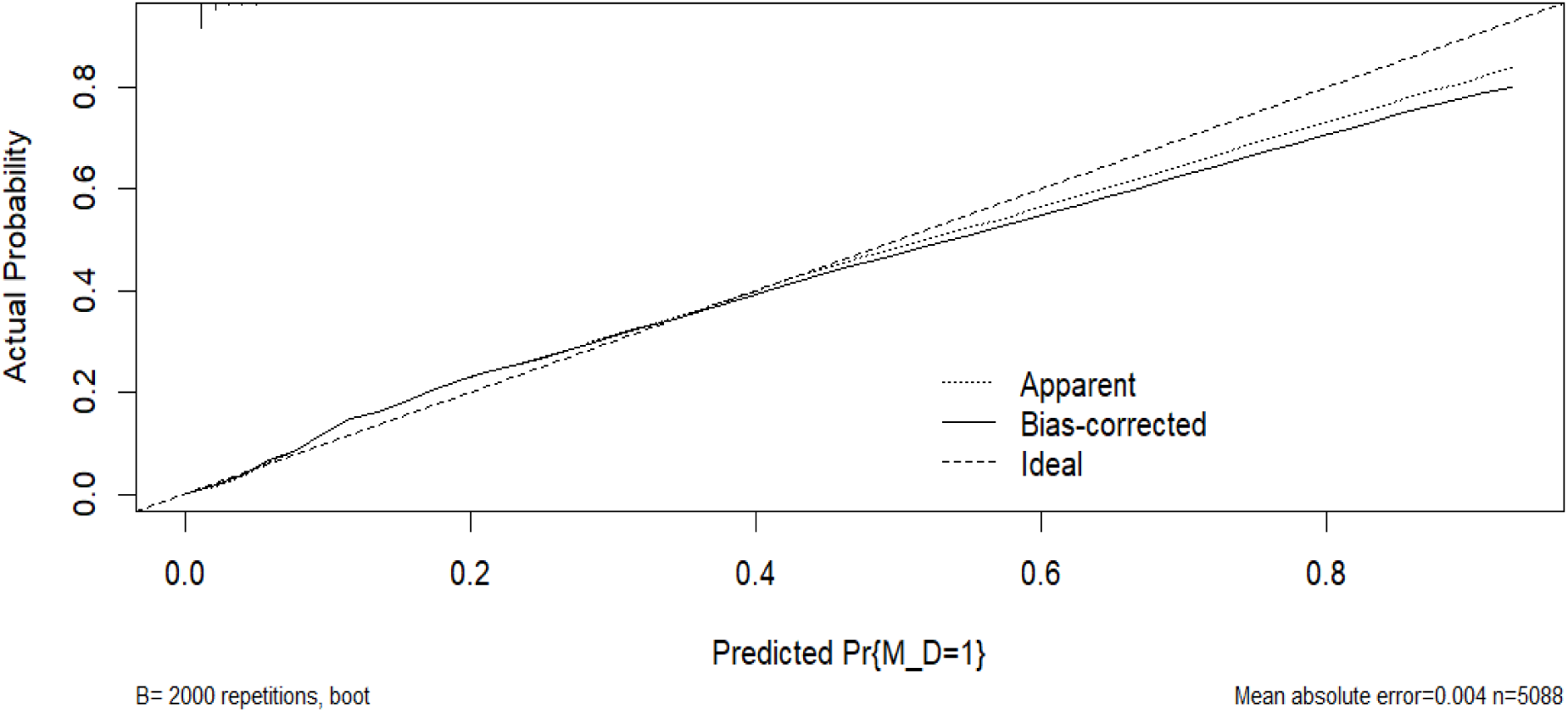
Calibration curve of validation set

**Fig 8.**
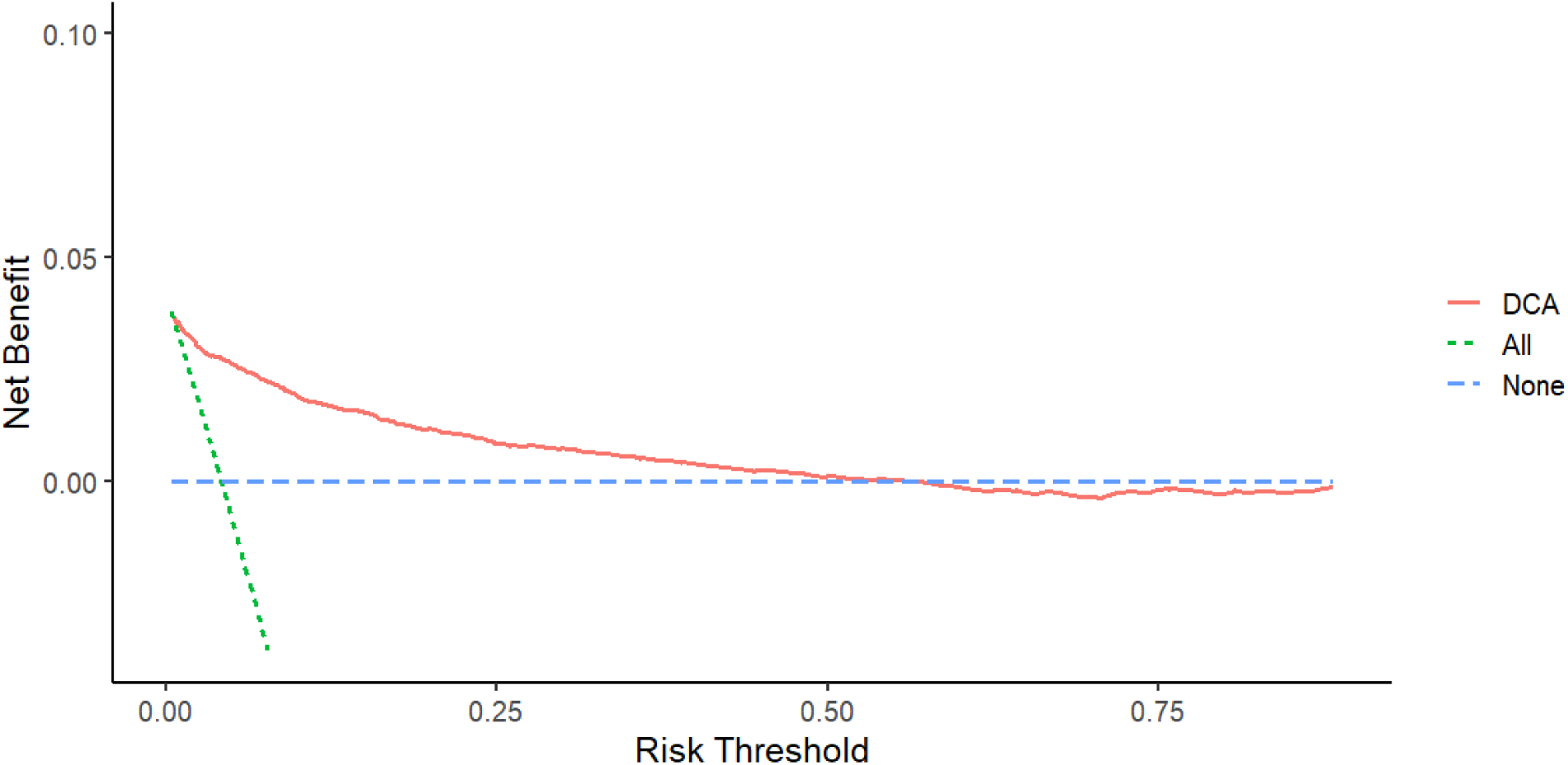
DCA curve of training set

**Fig 9.**
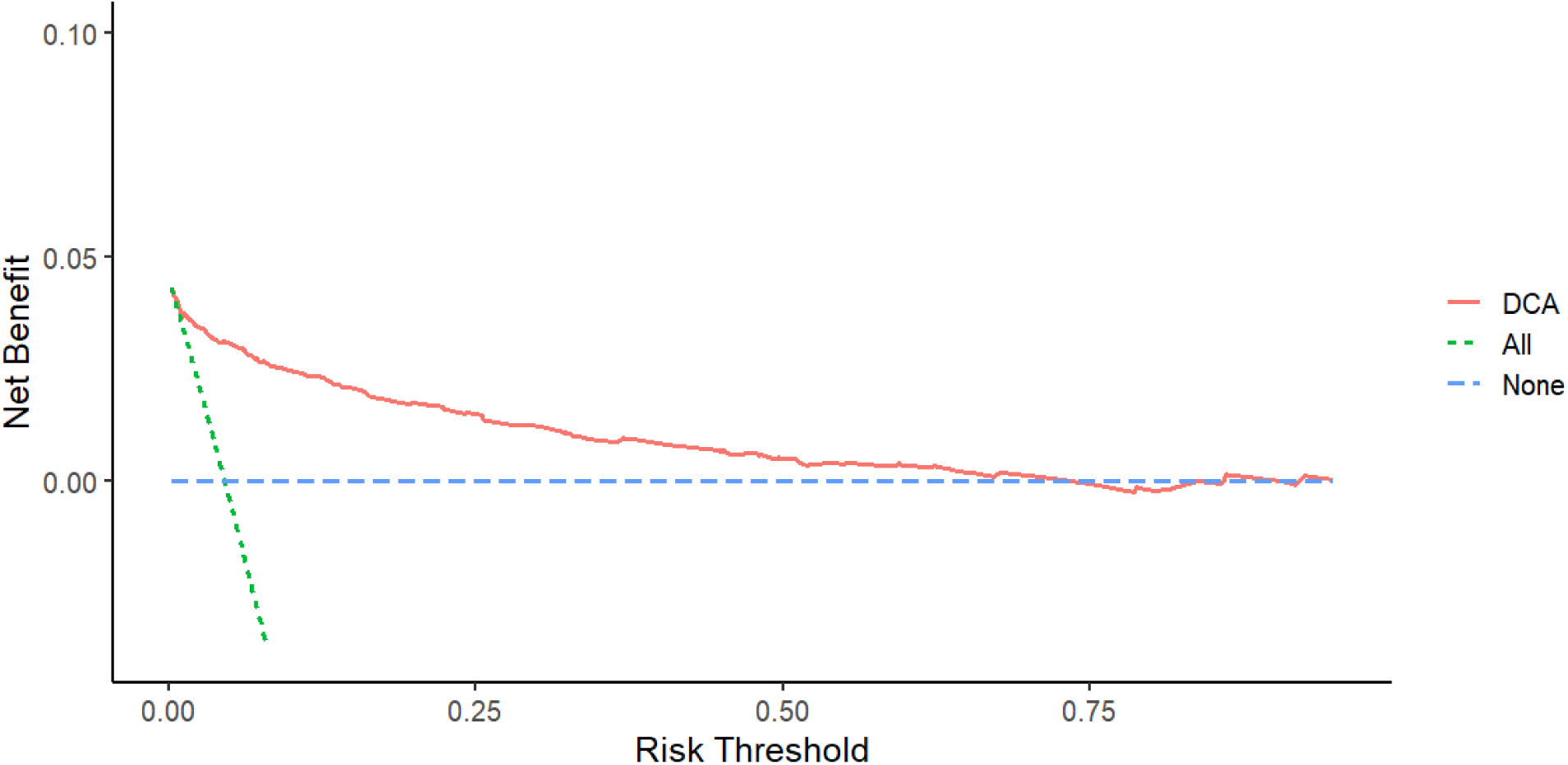
DCA Curve of Validation Set

### Prognostic factors of TNBC with distant metastasis

In this study, a cohort of 730 eligible patients with distant metastasis was assembled to investigate prognostic factors. As shown in Table 3.surgical intervention was undergone by 372 patients (51.0%), radiotherapy was received by 262 (35.9%), and chemotherapy was administered to 608 patients (83.3%). Both the chi-square test and Fisher’s exact test demonstrated no statistically significant differences across all variables between the training and validation sets (P > 0.05). Furthermore, univariate and multivariate prognostic regression analysis revealed that being unmarried (P < 0.001), not undergoing surgery (P < 0.001), and not receiving chemotherapy (P < 0.001) emerged as independent prognostic factors for TNBC patients with distant metastasis, as outlined in Table 4.

**Table 3.** Baseline clinical characteristics of patients diagnosed with TNBC accompanied by.

| | Training<br>(N=500) | Validation<br>(N=230) | $\chi^2$ | $P$ | Overall<br>(N=730) |
| --- | --- | --- | --- | --- | --- |
| **AgeGroup** |  |  | 0.462 | 0.793 |  |
| >60 | 206 (41.2%) | 98 (42.6%) |  |  | 304 (41.6%) |
| 20-44 | 102 (20.4%) | 42 (18.3%) |  |  | 144 (19.7%) |
| 45-59 | 192 (38.4%) | 90 (39.1%) |  |  | 282 (38.6%) |
| **Race** |  |  | 0.629 | 0.889 |  |
| AI/AN | 5 (1.0%) | 3 (1.3%) |  |  | 8 (1.1%) |
| Asian | 35 (7.0%) | 18 (7.8%) |  |  | 53 (7.3%) |
| Black | 106 (21.2%) | 44 (19.1%) |  |  | 150 (20.5%) |
| White | 354 (70.8%) | 165 (71.7%) |  |  | 519 (71.1%) |
| **Marital** |  |  | 0.003 | 0.959 |  |
| Married | 346 (69.2%) | 158 (68.7%) |  |  | 504 (69.0%) |
| Not Married | 154 (30.8%) | 72 (31.3%) |  |  | 226 (31.0%) |
| **Histology** |  |  | 0.539 | 0.763 |  |
| IDC | 406 (81.2%) | 188 (81.7%) |  |  | 594 (81.4%) |
| ILC | 28 (5.6%) | 10 (4.3%) |  |  | 38 (5.2%) |
| Other | 66 (13.2%) | 32 (13.9%) |  |  | 98 (13.4%) |
| **Grade** |  |  | 0.299 | 0.861 |  |
| I | 6 (1.2%) | 3 (1.3%) |  |  | 9 (1.2%) |
| II | 75 (15.0%) | 38 (16.5%) |  |  | 113 (15.5%) |
| III-IV | 419 (83.8%) | 189 (82.2%) |  |  | 608 (83.3%) |
| **T** |  |  | 1.112 | 0.774 |  |
| T1 | 73 (14.6%) | 28 (12.2%) |  |  | 101 (13.8%) |
| T2 | 166 (33.2%) | 76 (33.0%) |  |  | 242 (33.2%) |
| T3 | 102 (20.4%) | 46 (20.0%) |  |  | 148 (20.3%) |
| T4 | 159 (31.8%) | 80 (34.8%) |  |  | 239 (32.7%) |
| **N** |  |  | 2.156 | 0.541 |  |
| N0 | 115 (23.0%) | 42 (18.3%) |  |  | 157 (21.5%) |
| N1 | 215 (43.0%) | 105 (45.7%) |  |  | 320 (43.8%) |
| N2 | 60 (12.0%) | 28 (12.2%) |  |  | 88 (12.1%) |
| N3 | 110 (22.0%) | 55 (23.9%) |  |  | 165 (22.6%) |
| **SurgeryPrim** |  |  | 3.150 | 0.207 |  |
| Breast-Conserving | 74 (14.8%) | 23 (10.0%) |  |  | 97 (13.3%) |
| mastectomy | 185 (37.0%) | 90 (39.1%) |  |  | 275 (37.7%) |
| No | 241 (48.2%) | 117 (50.9%) |  |  | 358 (49.0%) |
| **NonPrimSurg** |  |  | 0.916 | 0.338 |  |
| No | 466 (93.2%) | 209 (90.9%) |  |  | 675 (92.5%) |
| Yes | 34 (6.8%) | 21 (9.1%) |  |  | 55 (7.5%) |
| **Radiation** |  |  | 2.258 | 0.132 |  |
| No | 311 (62.2%) | 157 (68.3%) |  |  | 468 (64.1%) |
| Yes | 189 (37.8%) | 73 (31.7%) |  |  | 262 (35.9%) |
| **Chemotherapy** |  |  | 0.193 | 0.659 |  |
| No | 81 (16.2%) | 41 (17.8%) |  |  | 122 (16.7%) |
| Yes | 419 (83.8%) | 189 (82.2%) |  |  | 608 (83.3%) |
| **TumorSize** |  |  | 3.088 | 0.213 |  |
| <50 | 301 (60.2%) | 123 (53.5%) |  |  | 424 (58.1%) |
| >100 | 48 (9.6%) | 28 (12.2%) |  |  | 76 (10.4%) |
| 50~100 | 151 (30.2%) | 79 (34.3%) |  |  | 230 (31.5%) |

**Table 4.**
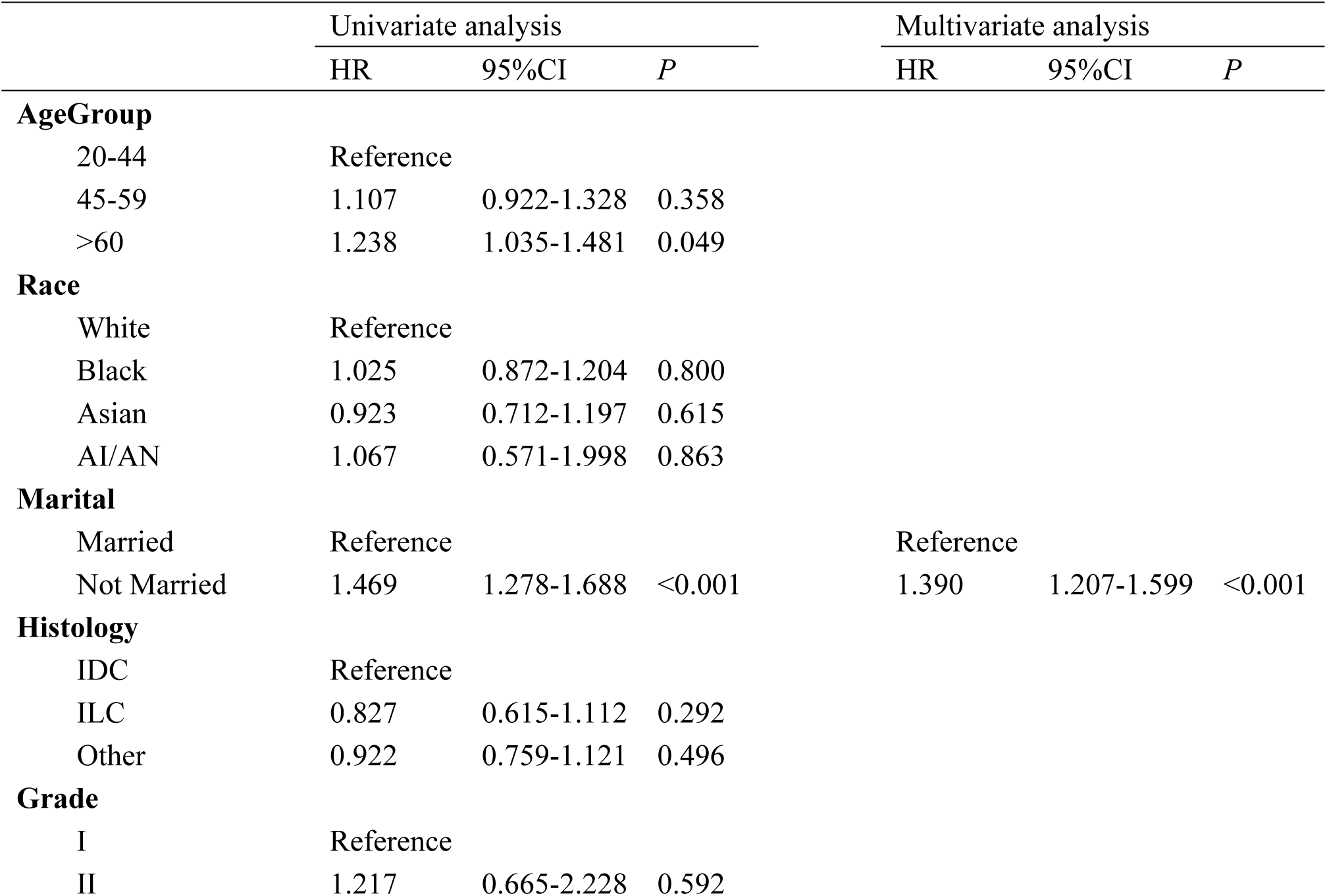
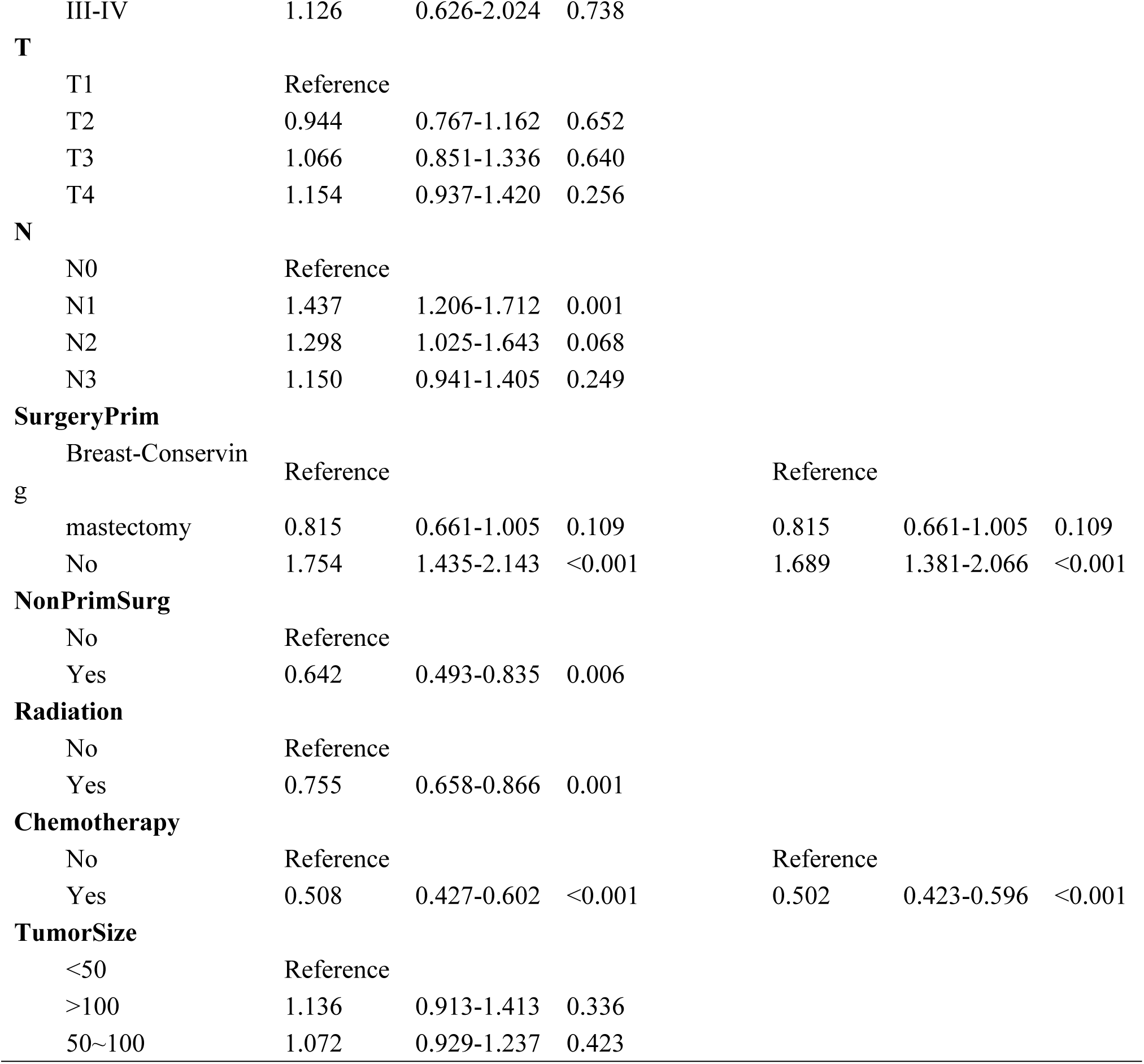
Univariate and multivariate Cox analysis of patients with distant metastasis of TNBC.

### Prognostic nomogram and validation

Based on these three prognostic factors, we constructed a nomogram to predict the OS of patients with distant metastases (Fig 10). Calibration curves were plotted separately in the training set (Figs 11-13) and validation set (Figs 14-16), showing strong consistency between the predicted OS probabilities at 12, 36, and 60 months and the actual outcomes. The nomogram’s good performance in clinical practice was determined by DCA (Figs 17-21). The ROC in the training set showed AUCs of 0.751, 0.709, and 0.691 at 12, 36, and 60 months, respectively, while the ROC in the validation set showed AUCs of 0.690, 0.690, and 0.732 (Figs 22 and 23).

**Fig 10.**
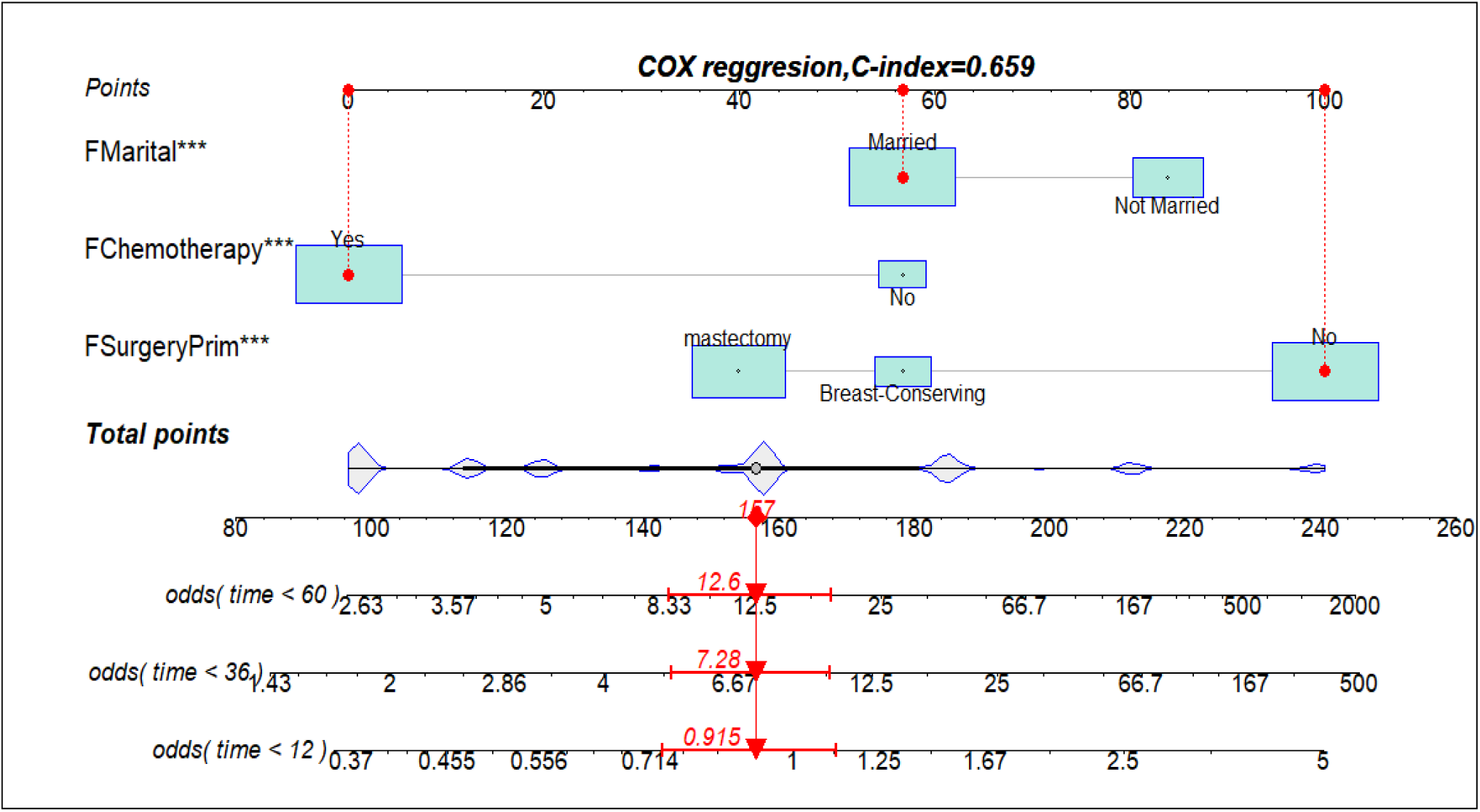
Prognostic nomogram for predicting OS in TNBC patients with distant metastasis at 12, 24, and 36 months

**Fig 11.**
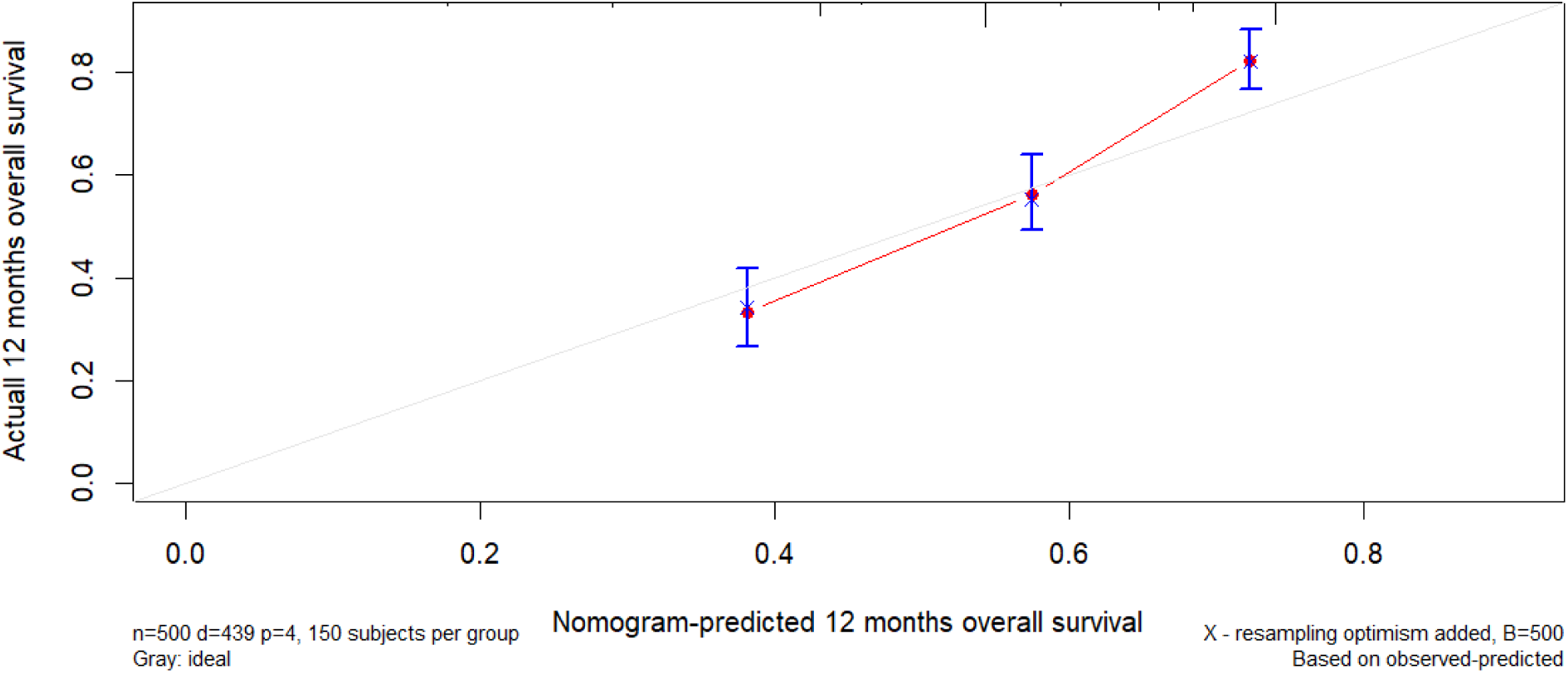
Calibration curve for the training set over 12 months

**Fig 12.**
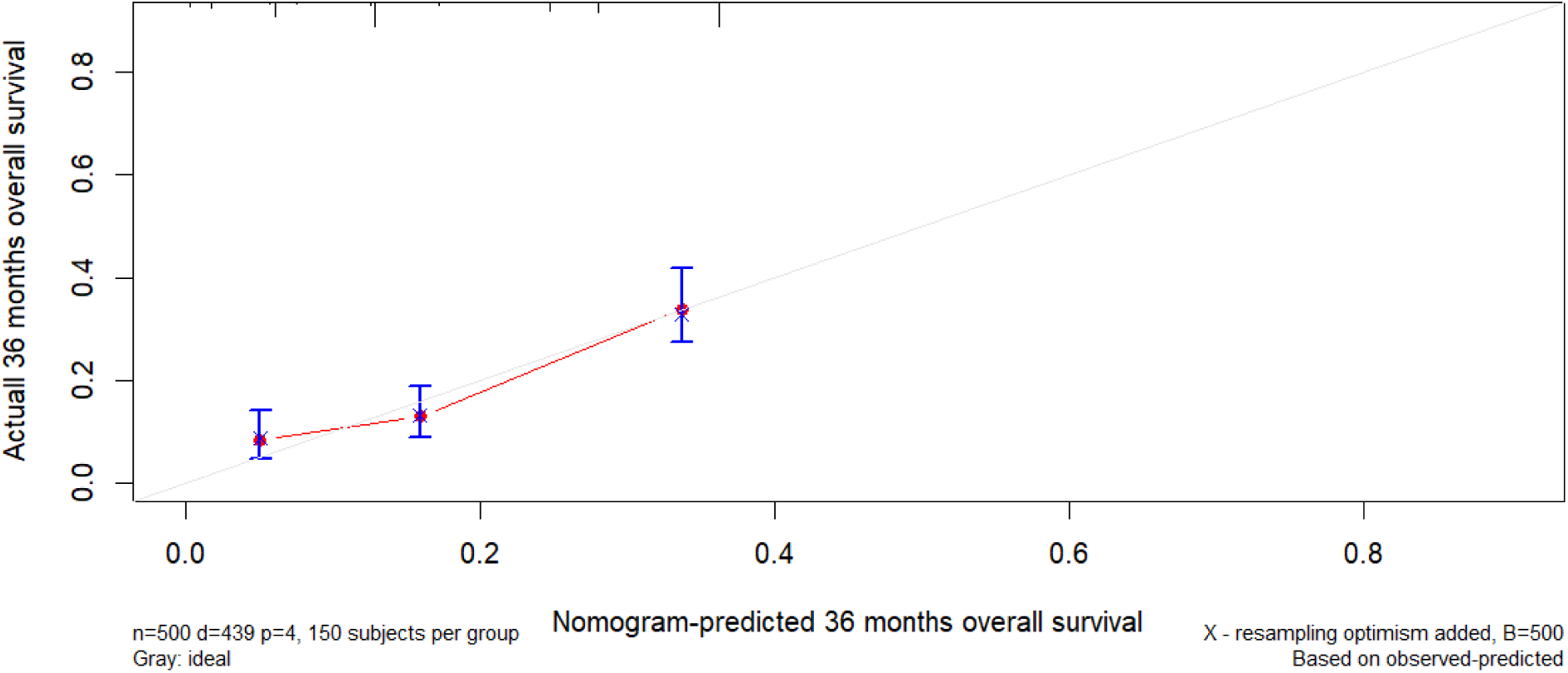
Calibration curve for the training set over 36 months

**Fig 13.**
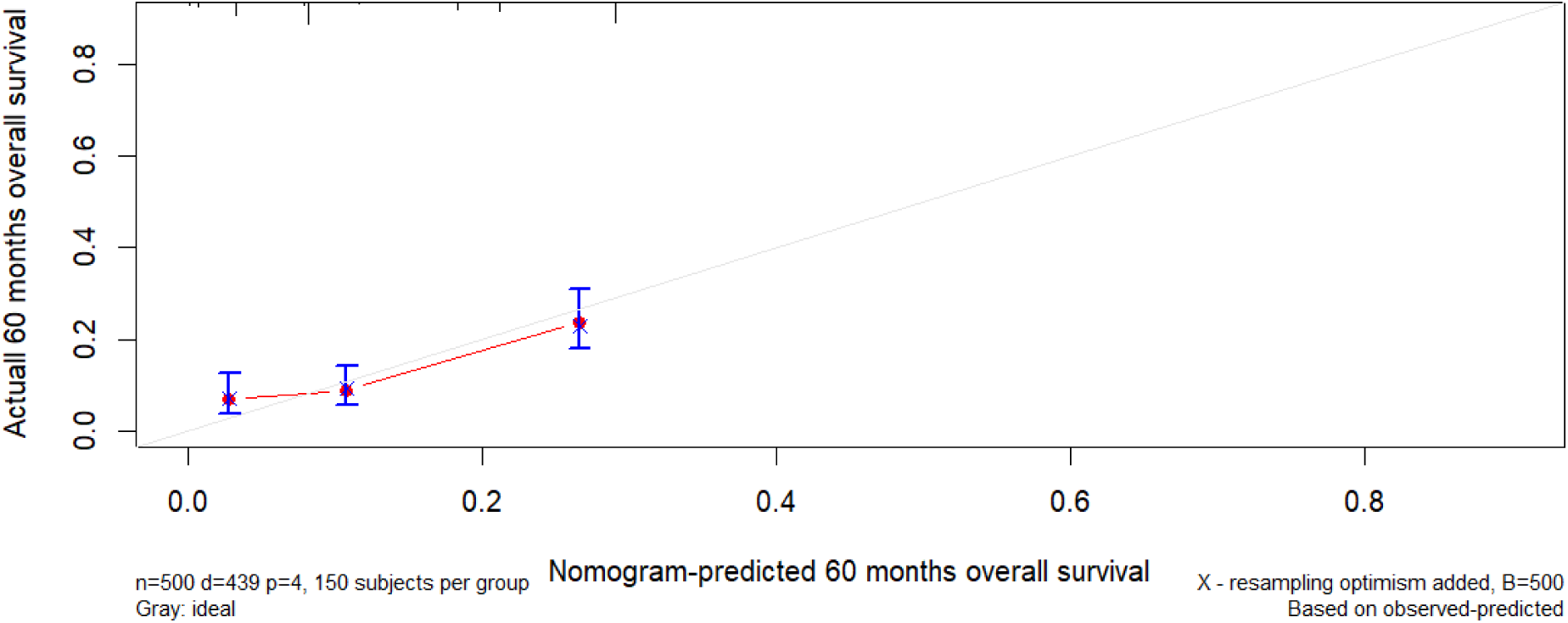
Calibration curve for the training set over 60 month

**Fig14.**
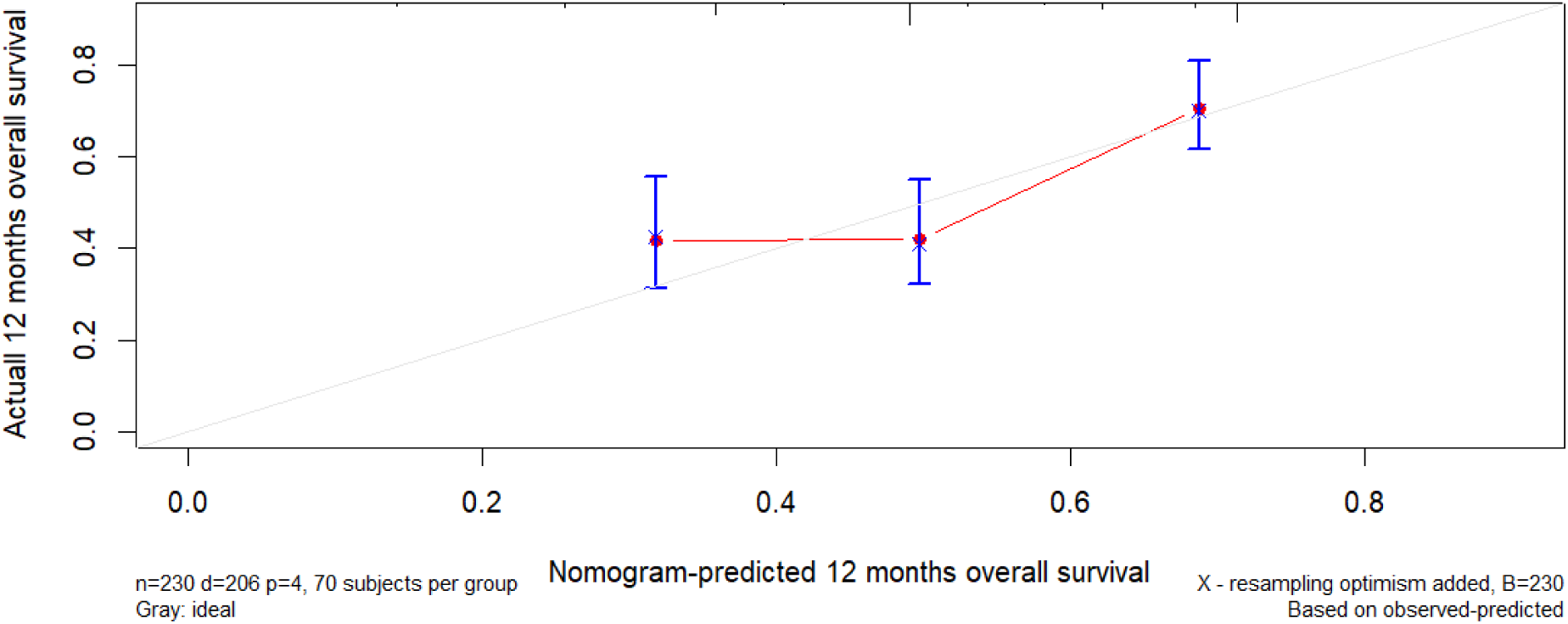
Calibration curve for the validation set over 12 months

**Fig 15.**
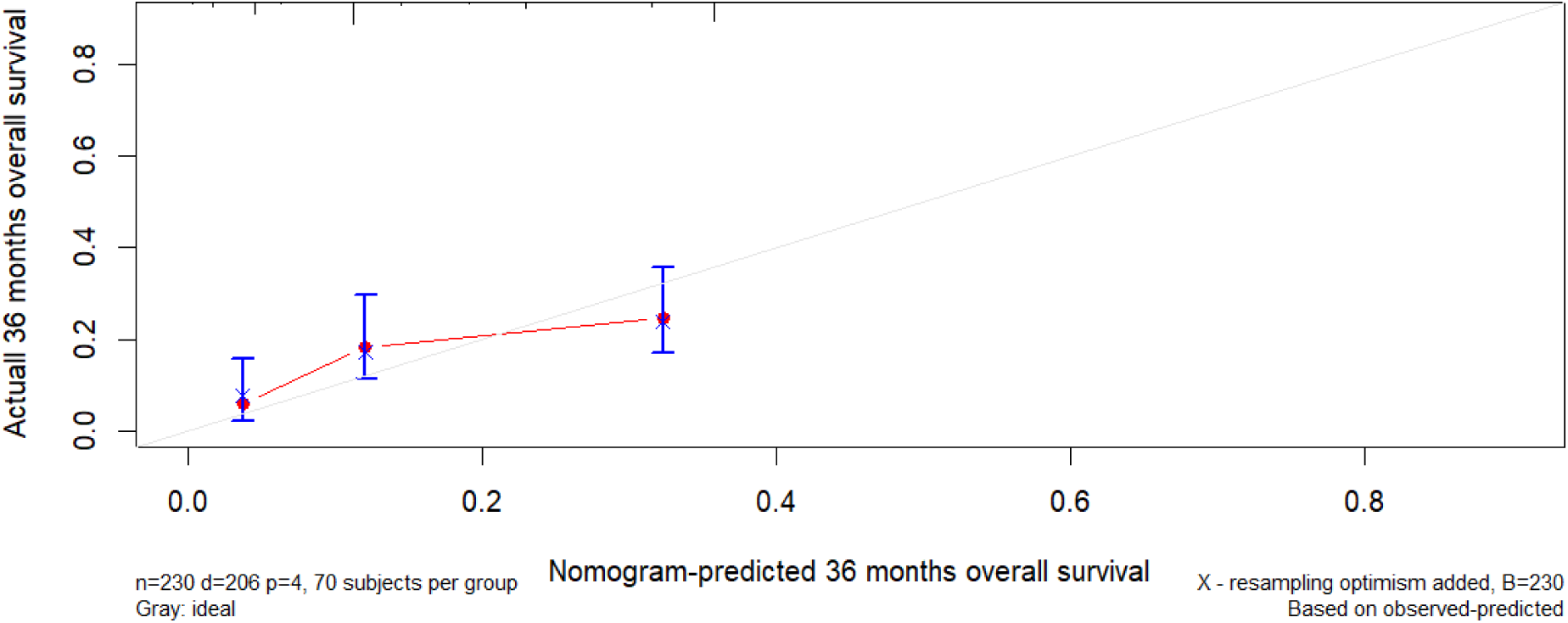
Calibration curve for the validation set over 36 months

**Fig 16.**
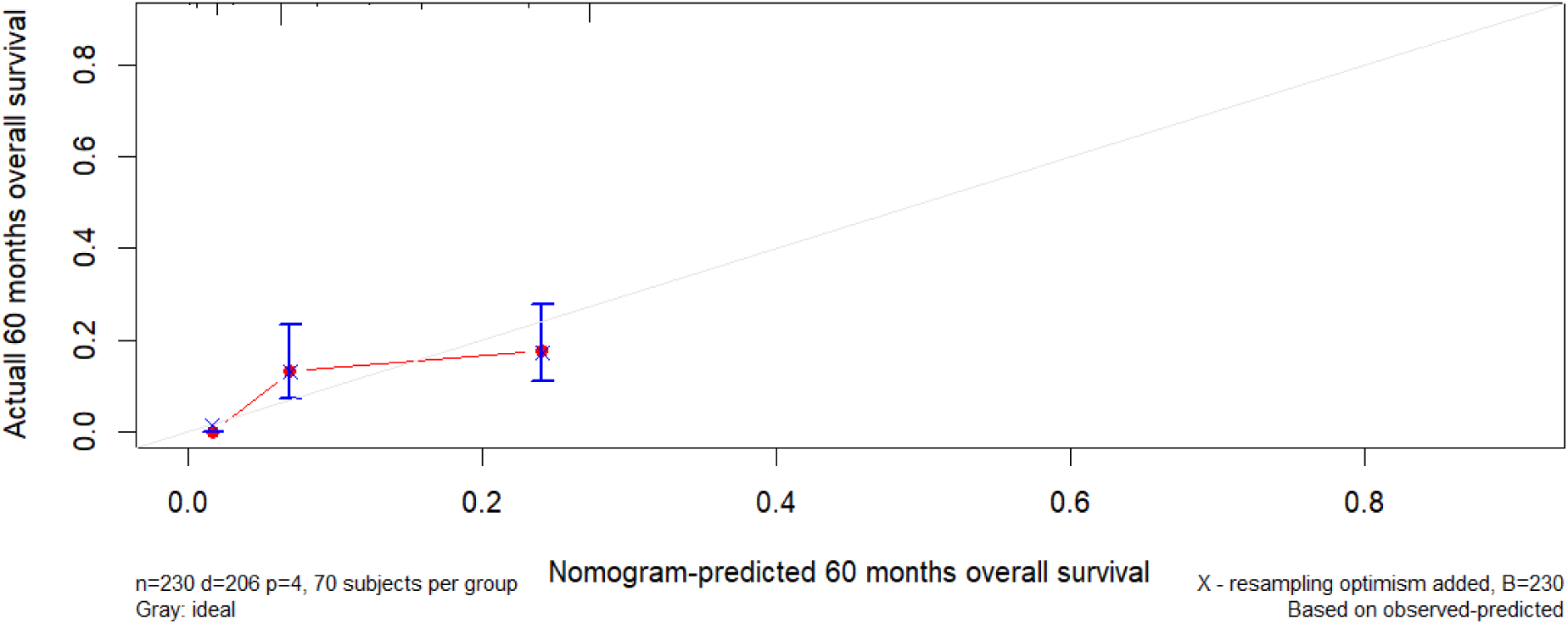
Calibration curve for the validation set over 60 months

**Fig 17.**
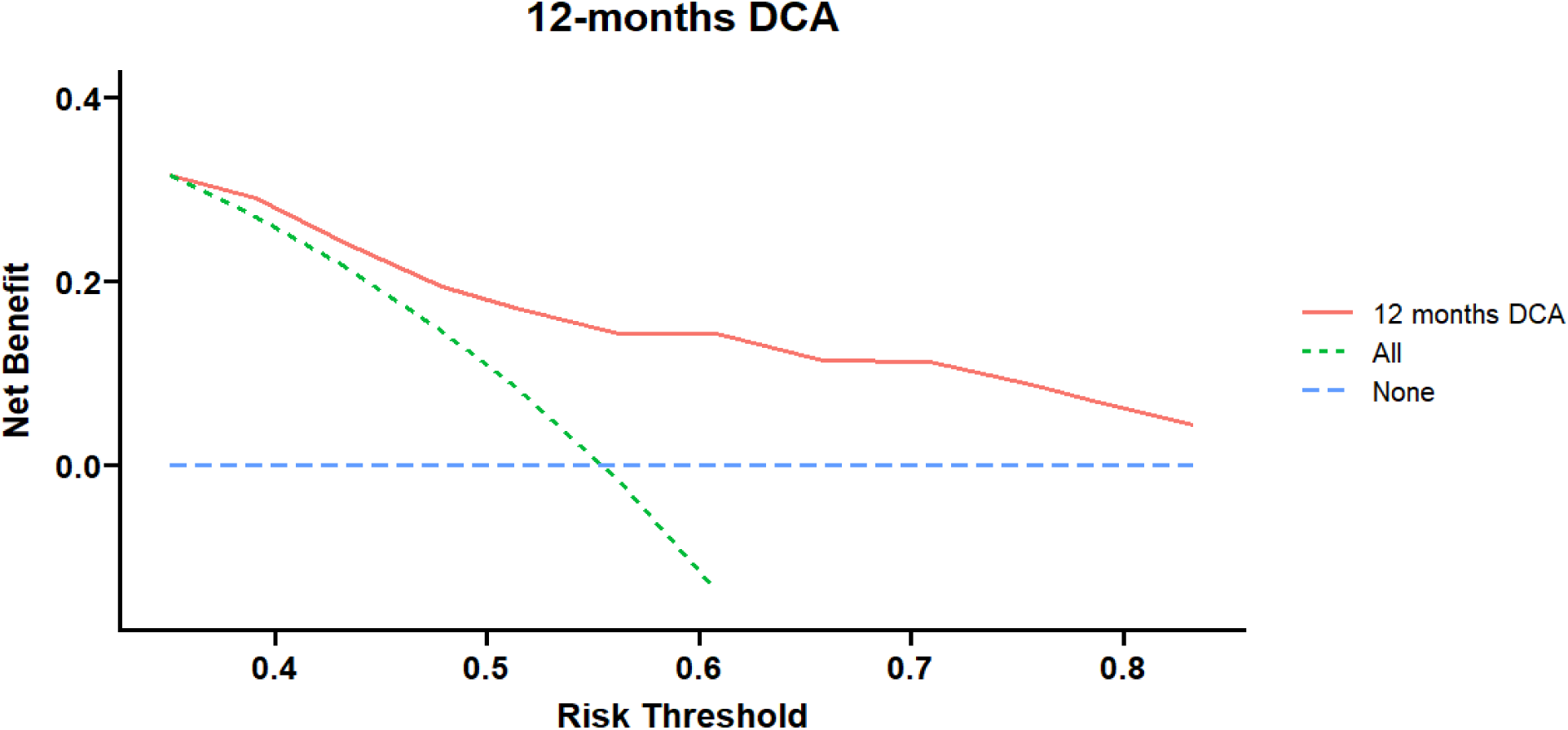
DCA Curve of the Training Set over 12 Months

**Fig 18.**
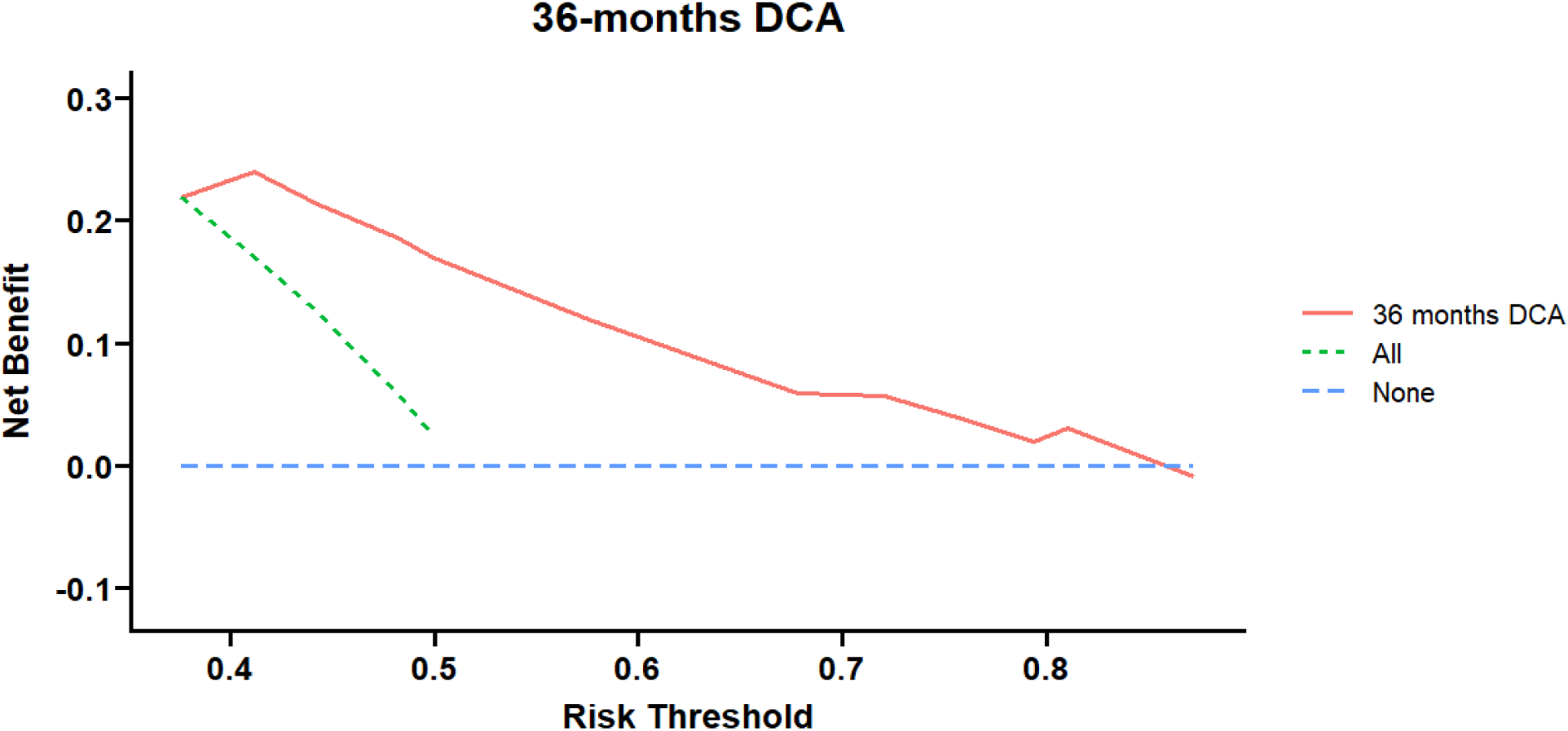
DCA Curve of the Training Set over 36 Months

**Fig 19.**
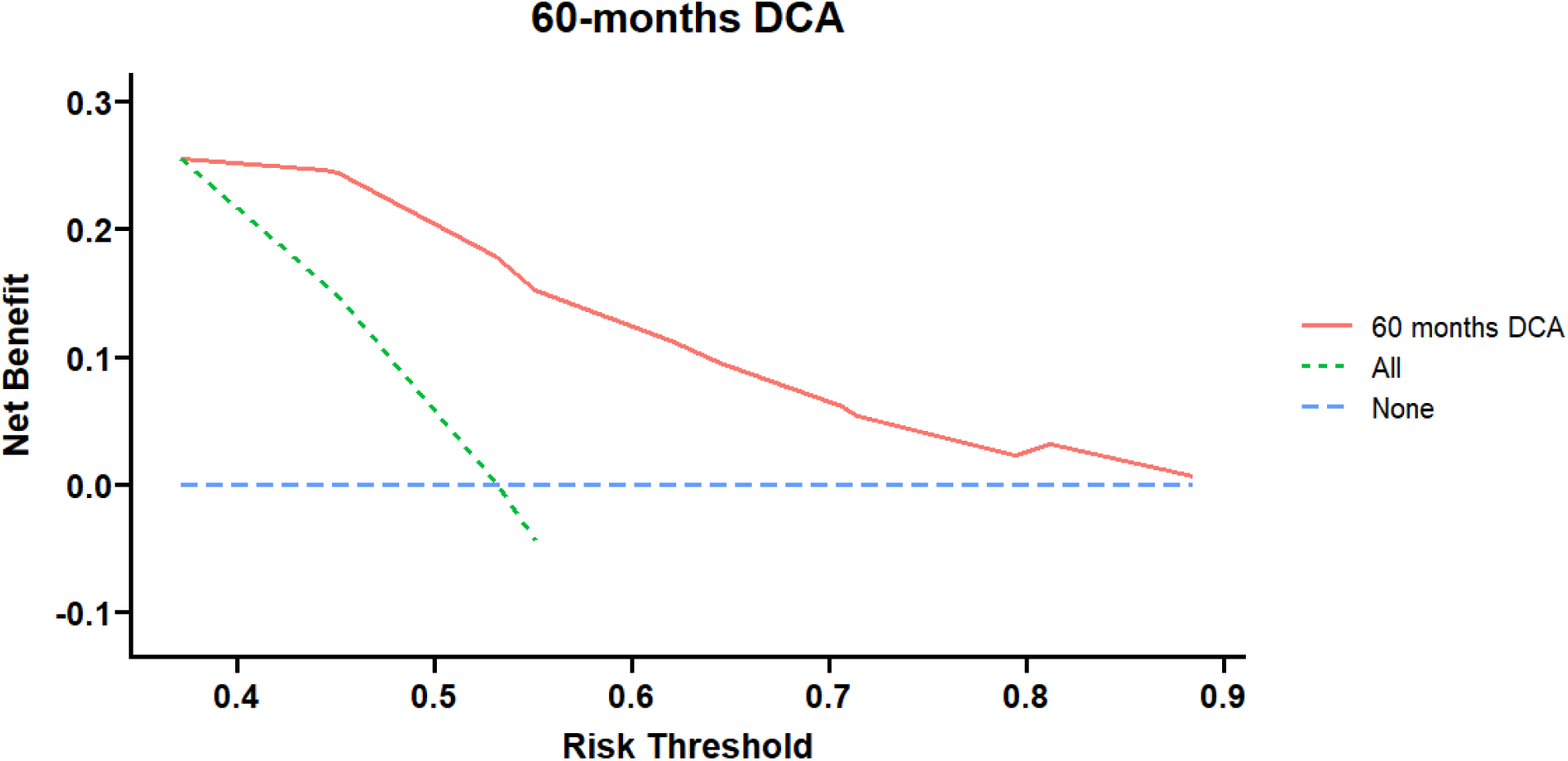
DCA Curve of the Training Set over 60 Months

**Fig 20.**
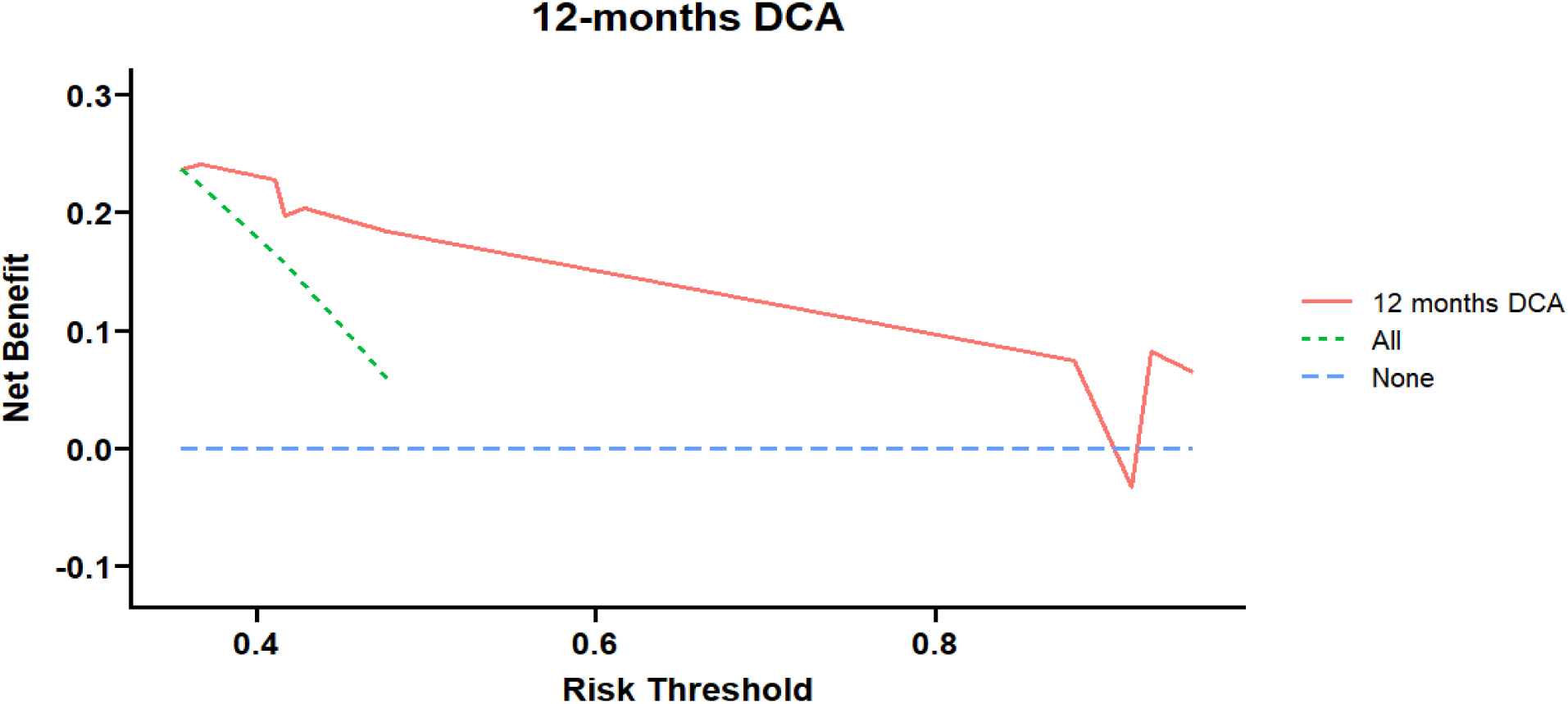
DCA Curve of the Validation Set over 12 Months

**Fig 21.**
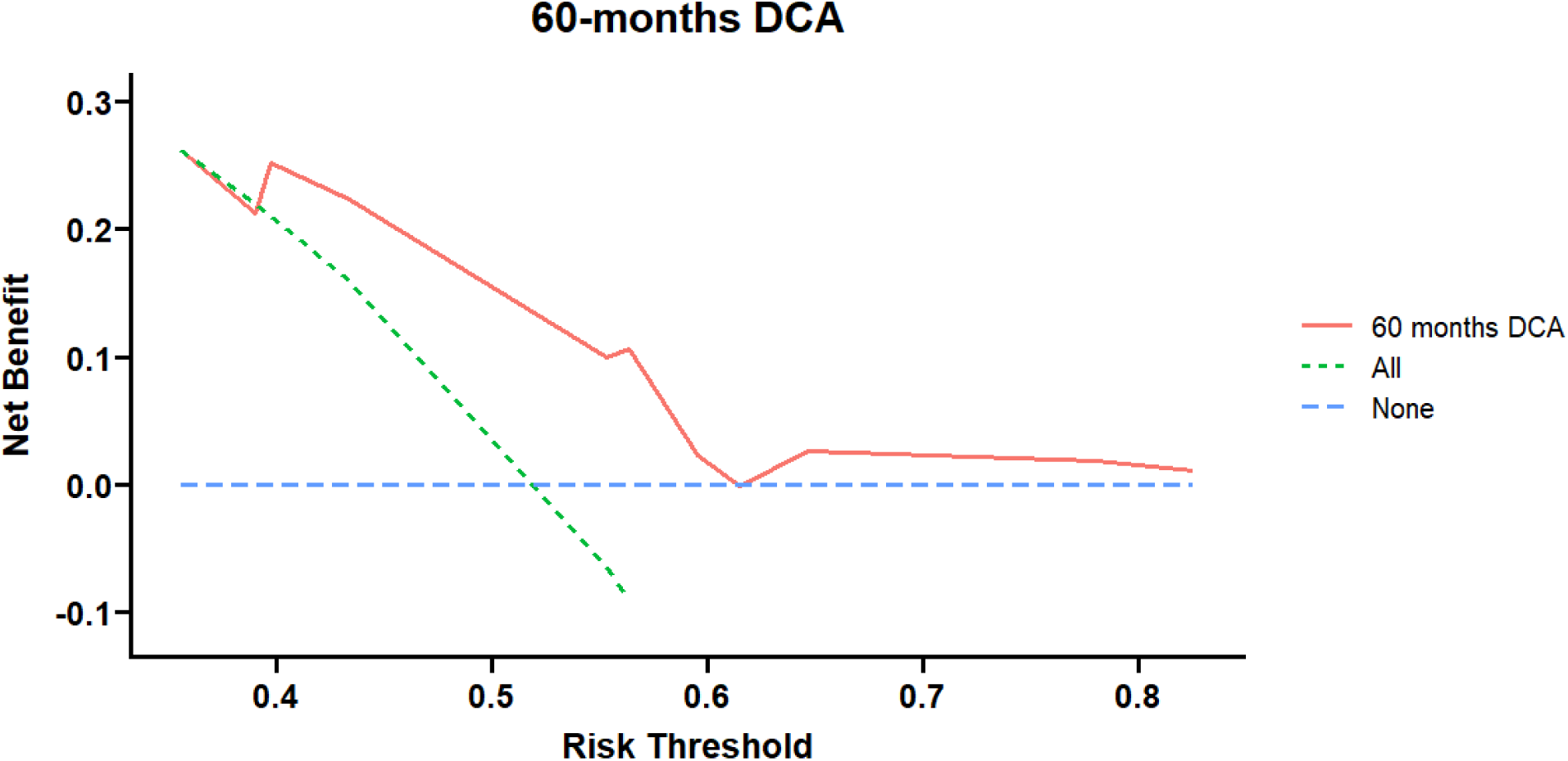
DCA Curve of the Validation Set over 60 Months

**Fig 22.**
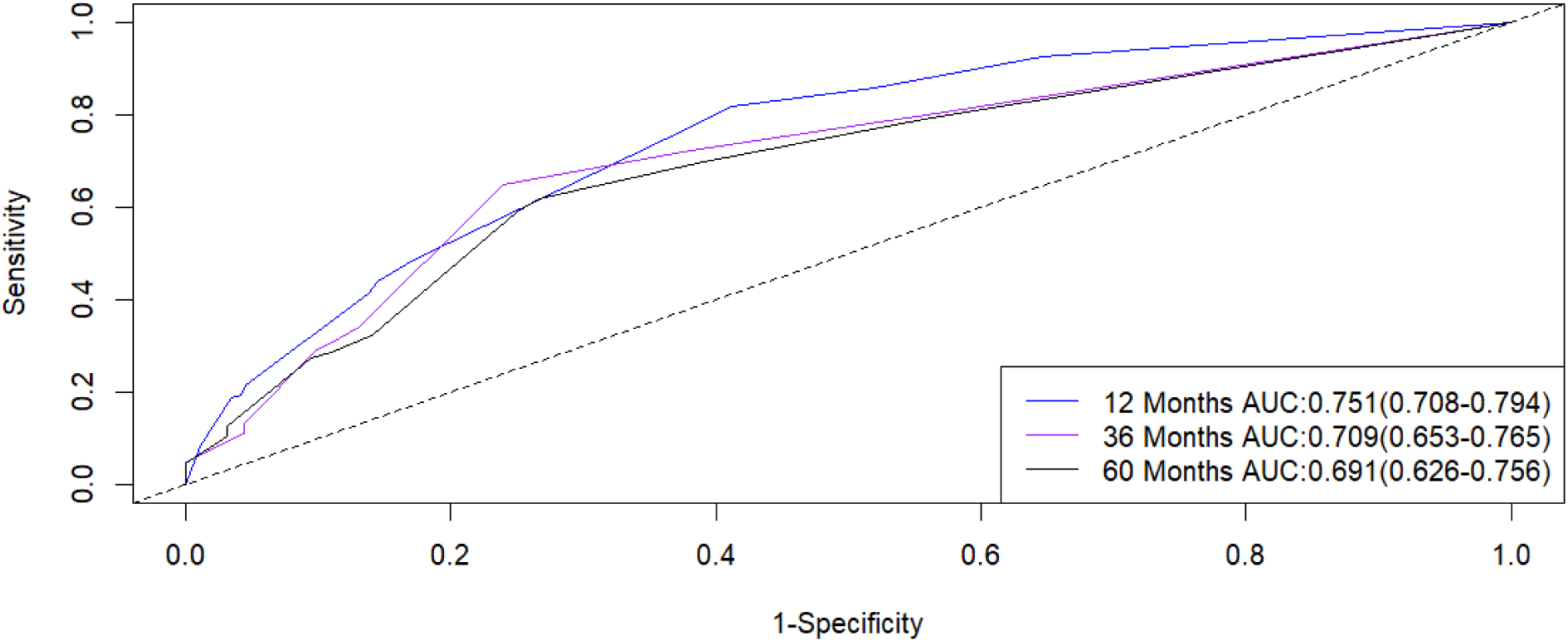
Receiver Operating Characteristic Curves for Subjects at 12, 36, and 60 Months in the Training Set

**Fig 23.**
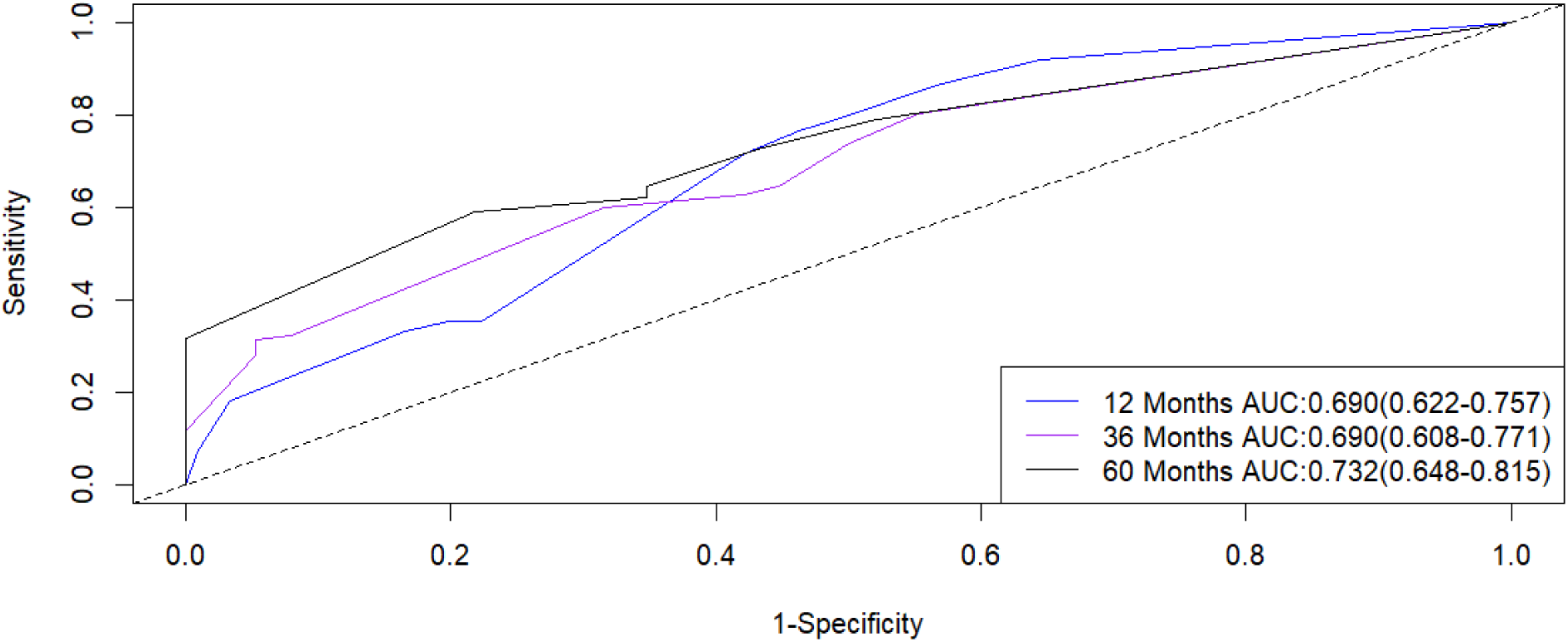
Receiver Operating Characteristic Curves for Subjects at 12, 36, and 60 Months in the Validation Set

Finally, each patient was assigned a risk score based on the three features of the nomogram, and a risk classification was developed based on this. The K-M curves showed that the OS of patients in the high-risk group was significantly lower than that of the low-risk group (Figs 24 and 25). At the same time, by comparing the nomogram with each independent prognostic factor at 12, 36, and 60 months, it was shown that the nomogram outperformed all independent prognostic factors (Figs 26-31).

**Fig 24.**
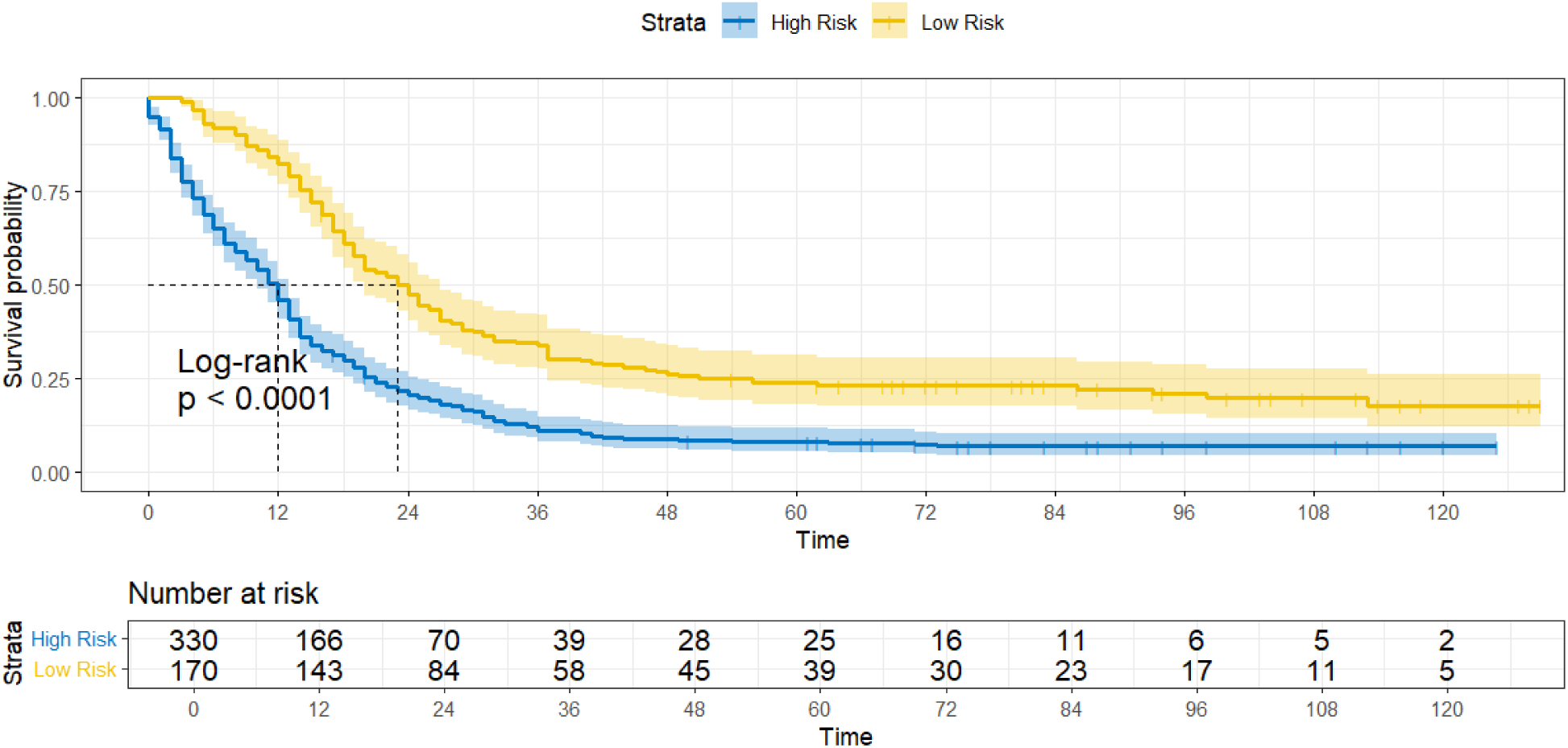
Overall Survival (OS) of high-risk and low-risk groups in the training set

**Fig 25.**
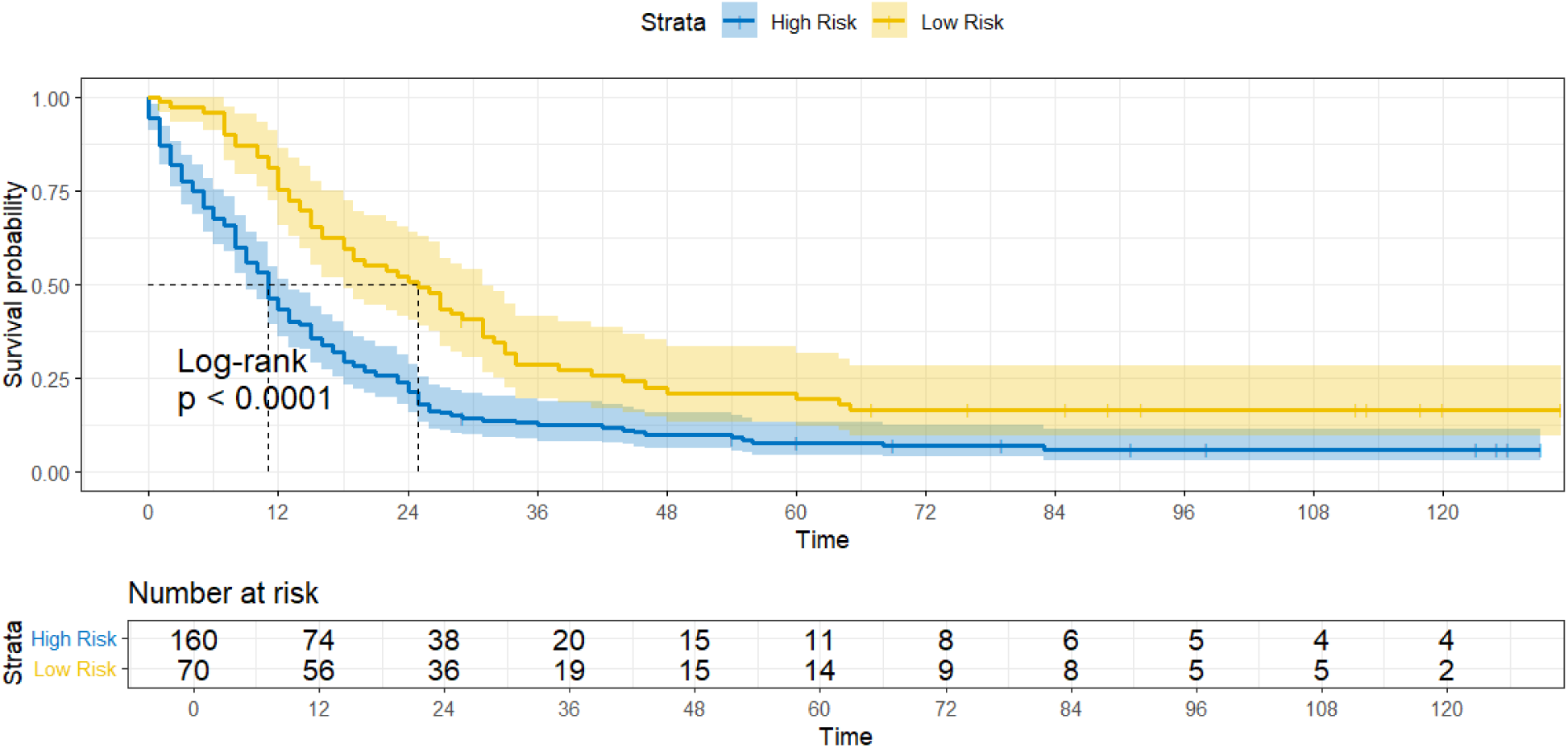
Overall Survival (OS) of high-risk and low-risk groups in the validation set

**Fig 26.**
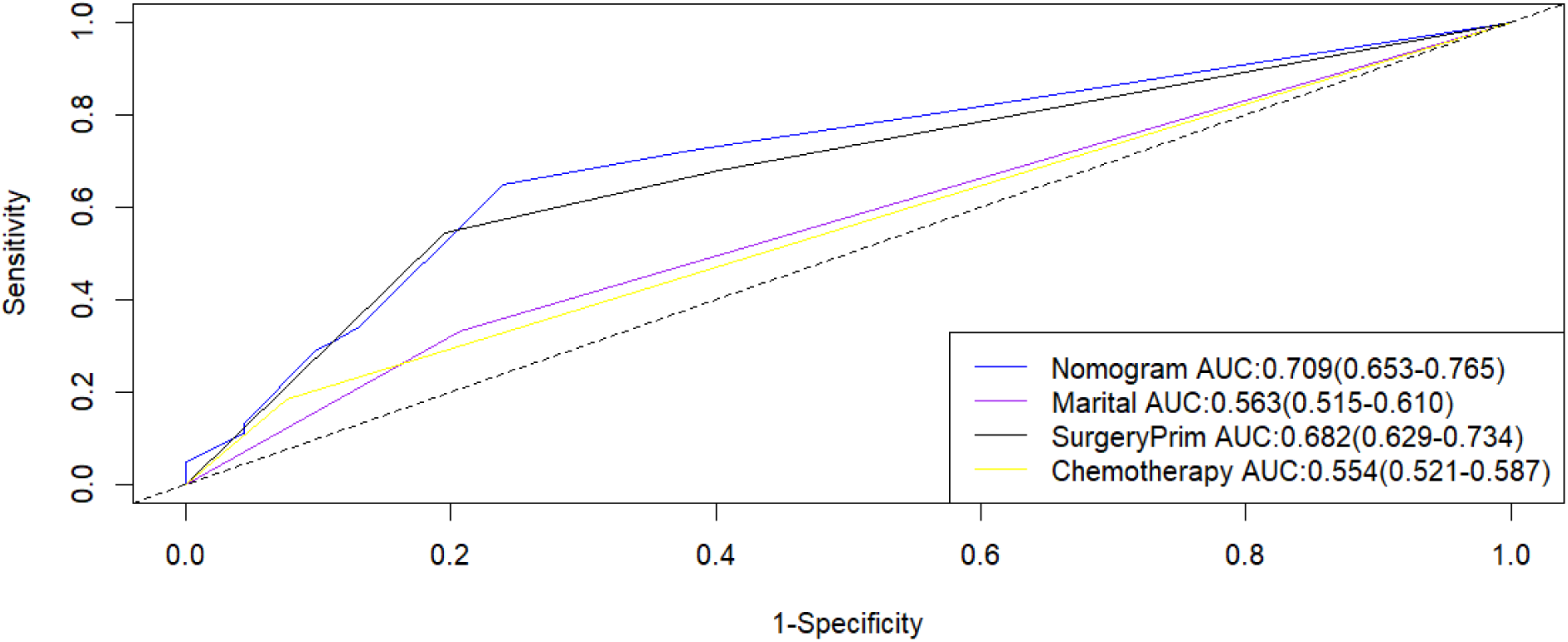
Comparison of the area under the receiver operating characteristic curve for the 12 month training set

**Fig 27.**
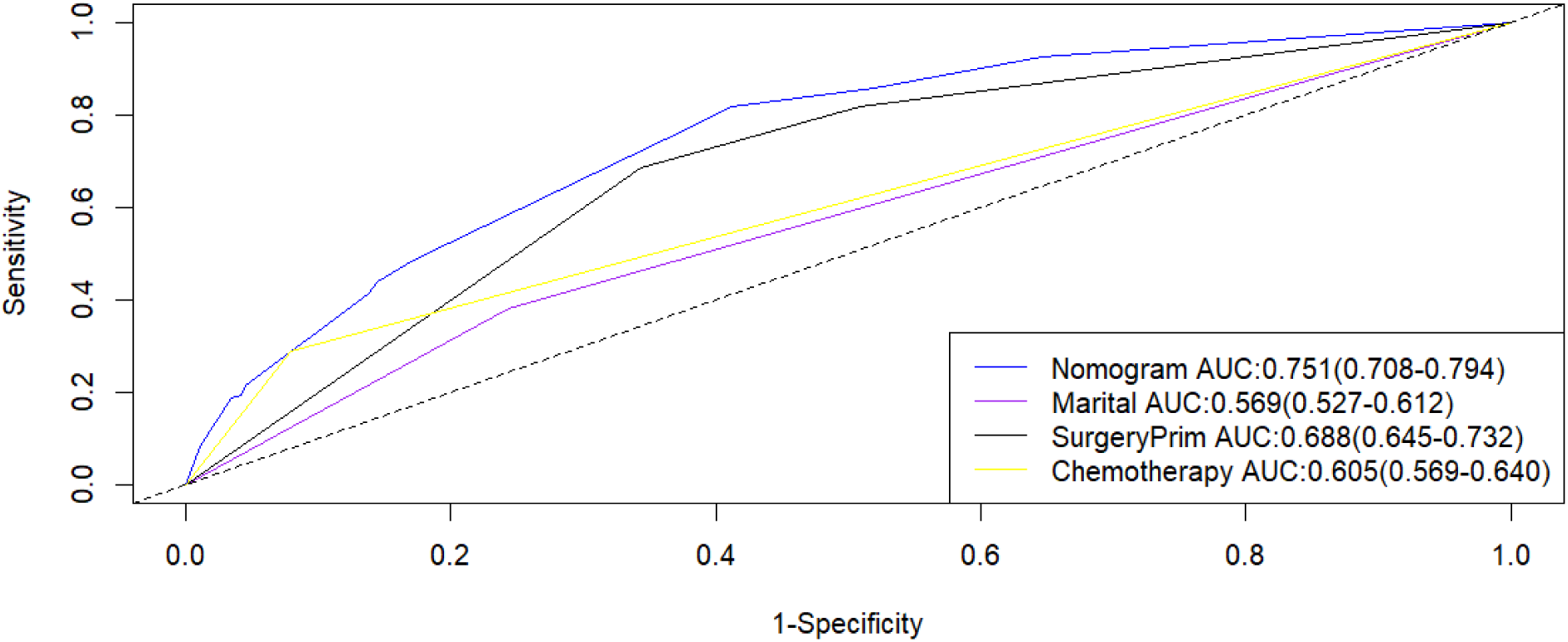
Comparison of the area under the receiver operating characteristic curve for the 36 month training set

**Fig 28.**
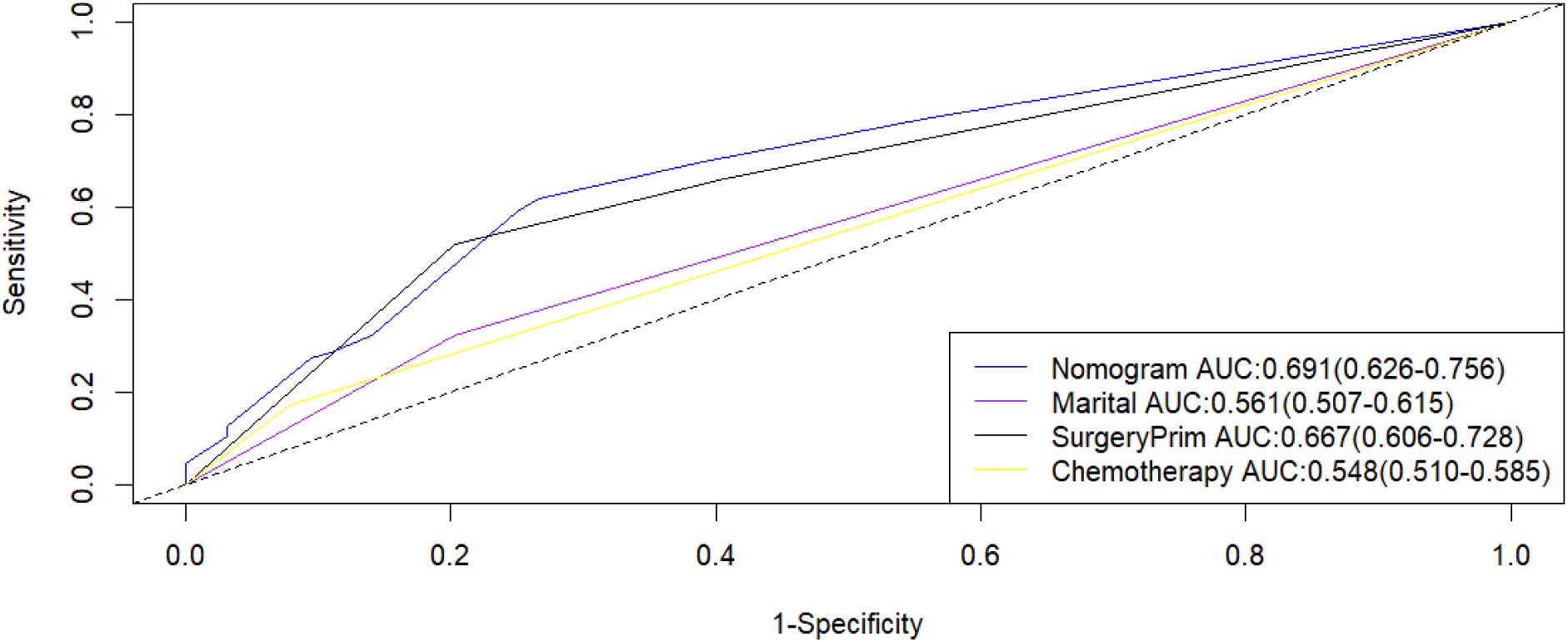
Comparison of the area under the receiver operating characteristic curve for the 60 month training set

**Fig 29.**
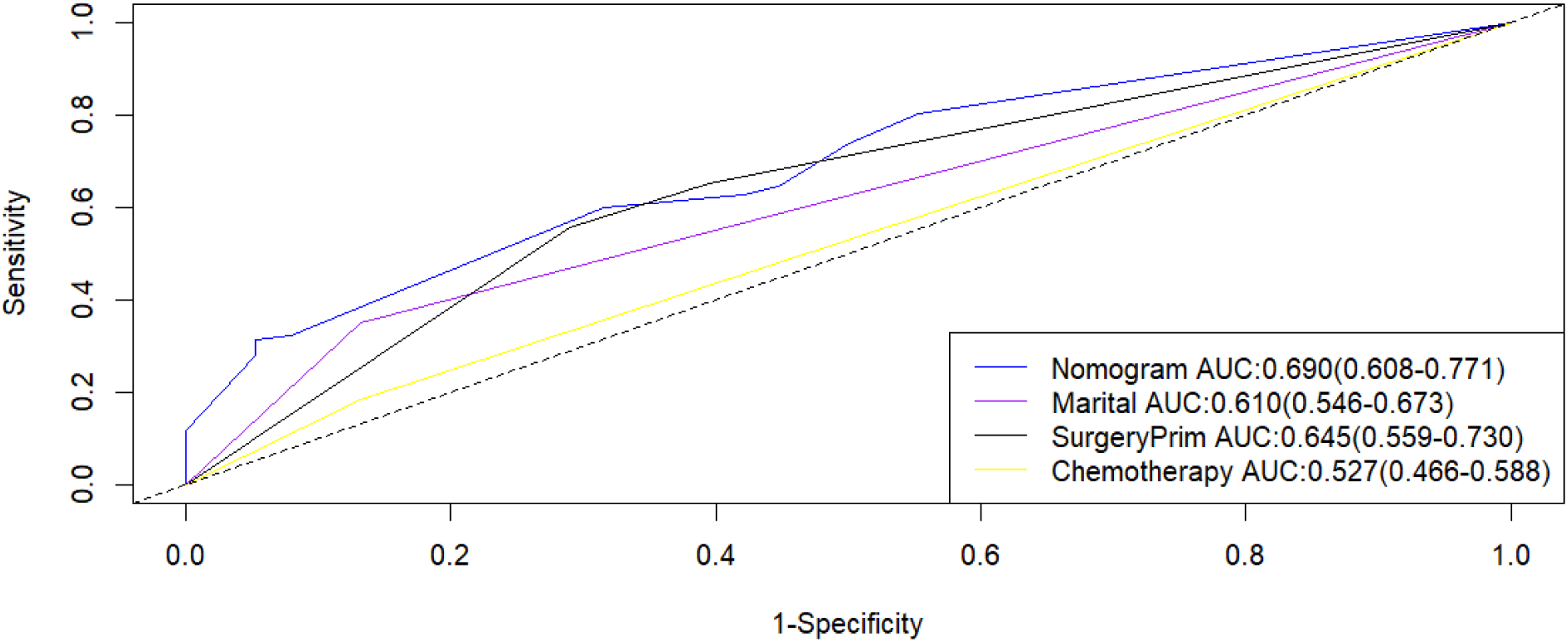
Comparison of the area under the receiver operating characteristic curve for the 12 month validation set

**Fig 30.**
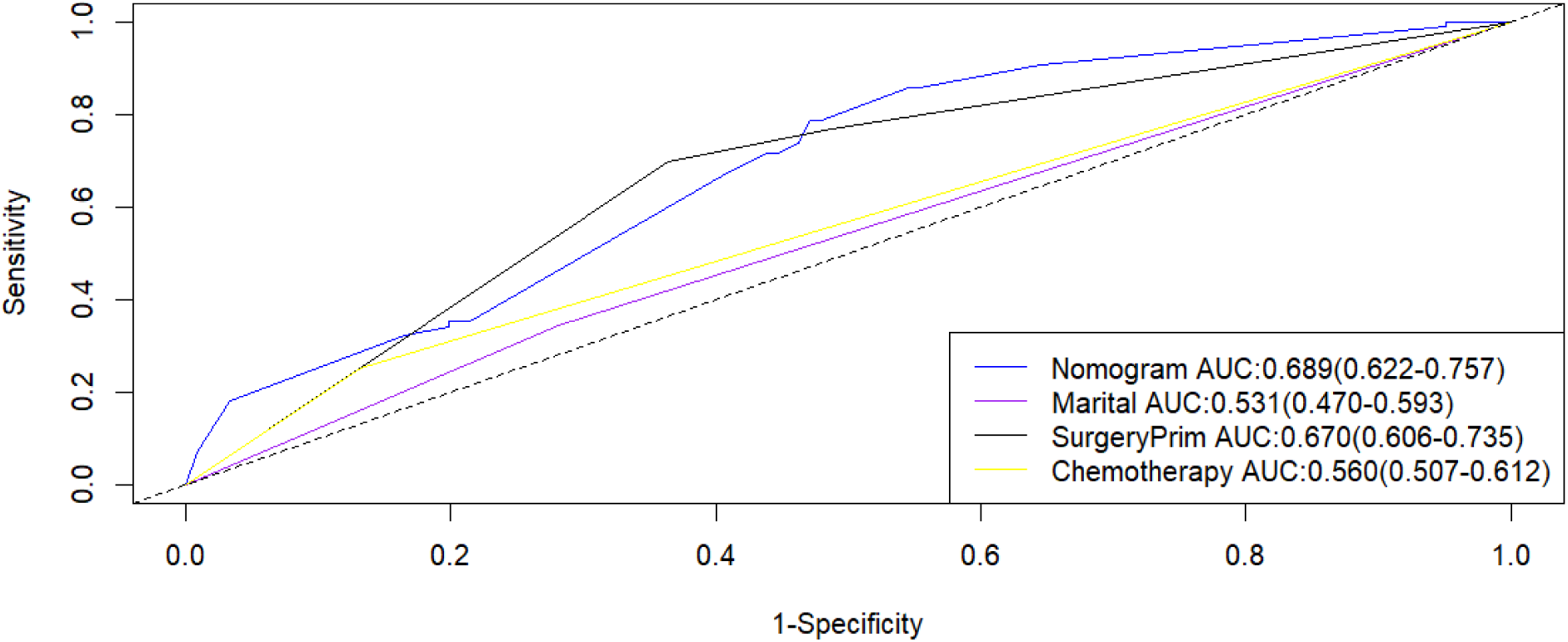
Comparison of the area under the receiver operating characteristic curve for the 36 month validation set

**Fig 31.**
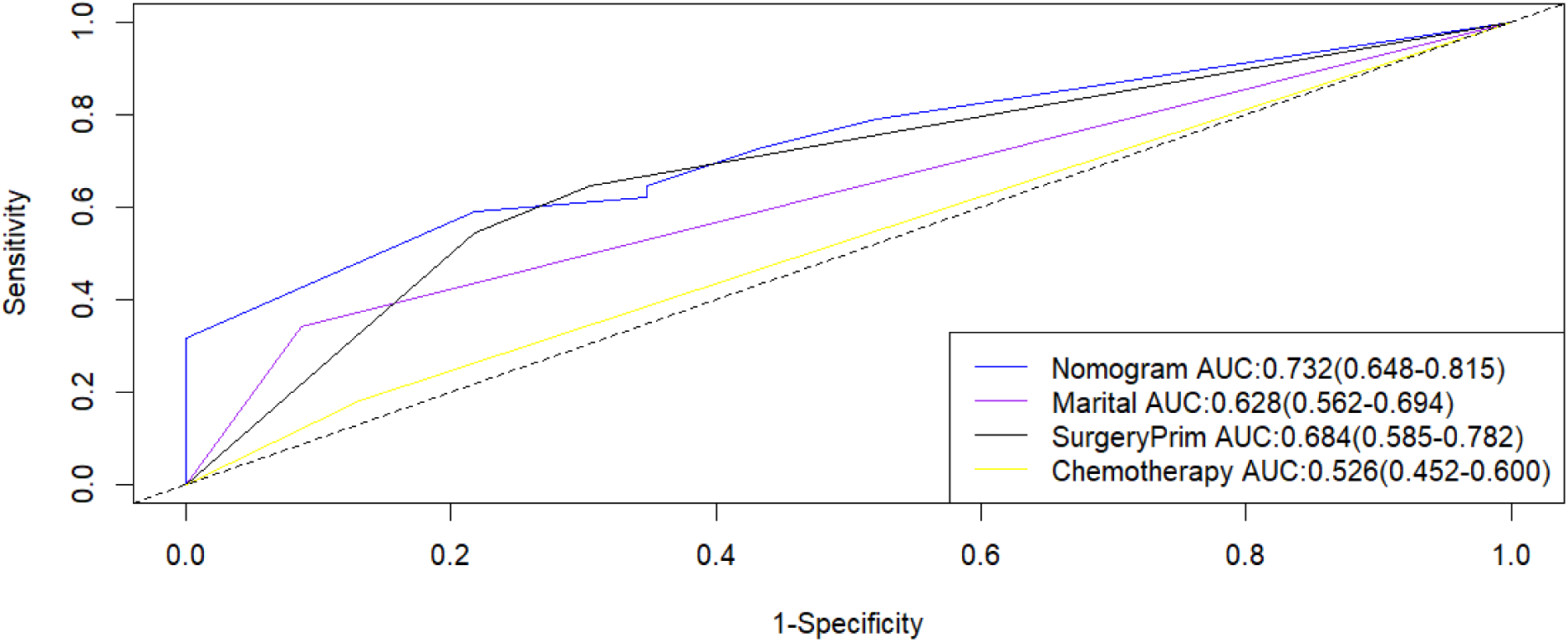
Comparison of the area under the receiver operating characteristic curve for the 60 month validation set

### Validating the performance of nomogram on an extended test set

An expanded test set was established by re-screening the SEER database to identify 157 TNBC patients diagnosed with distant metastasis in 2017, all having comprehensive information regarding marital status, chemotherapy, and surgery. Within this expanded test set, the calibration curve of the prognostic nomogram demonstrated a high degree of concordance between the predicted overall survival of patients with distant metastasis and the actual observed outcomes (Figs 32-34). Furthermore, the prognostic nomogram emerged as an effective clinical tool, as evidenced by the DCA evaluation (Figs 35-37). Notably, at 12, 36, and 60 months, the nomogram surpassed the performance of three independent predictors (Figs 38-40). Additionally, the AUCs for overall survival at 12, 36, and 60 months were 0.741, 0.727, and 0.718, respectively. The K-M survival analysis revealed significant differences in survival patterns between patients categorized as high-risk and low-risk (P<0.05) (Fig 41).

**Fig 32.**
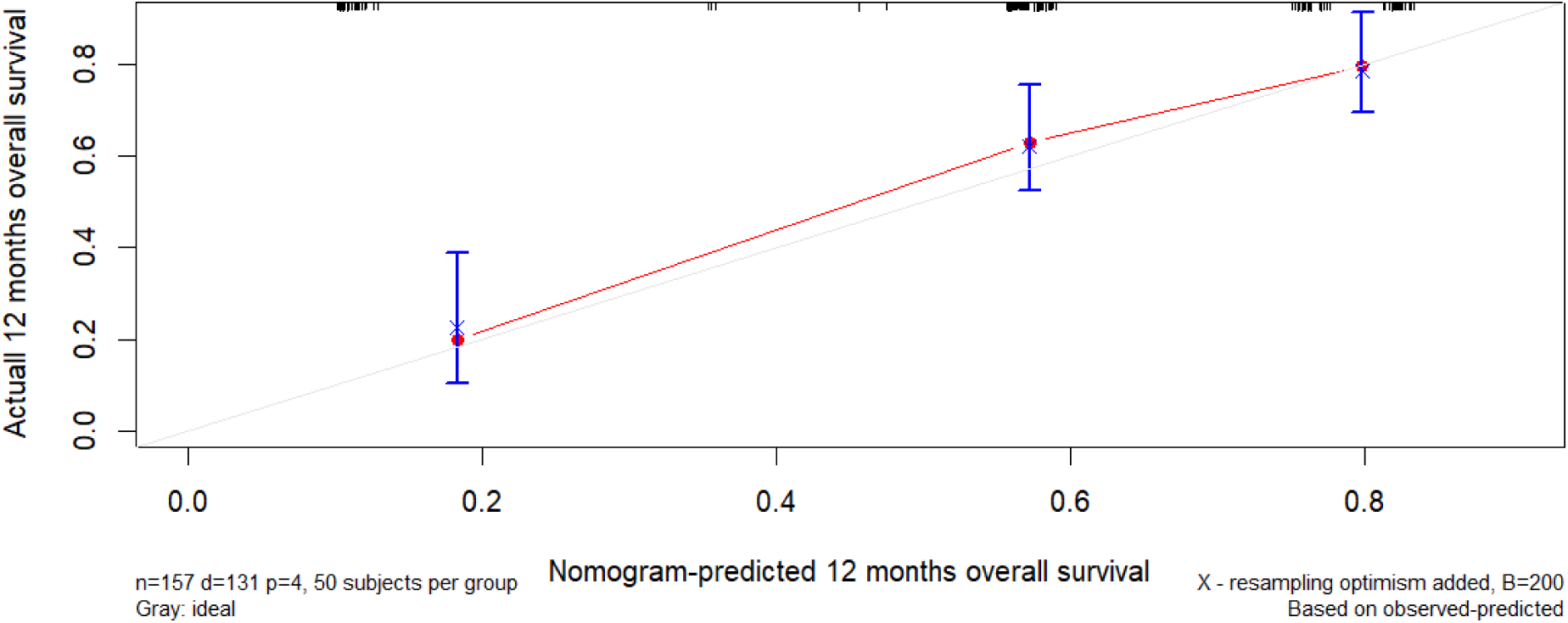
Calibration curve of the extended set 12 month nomogram

**Fig 33.**
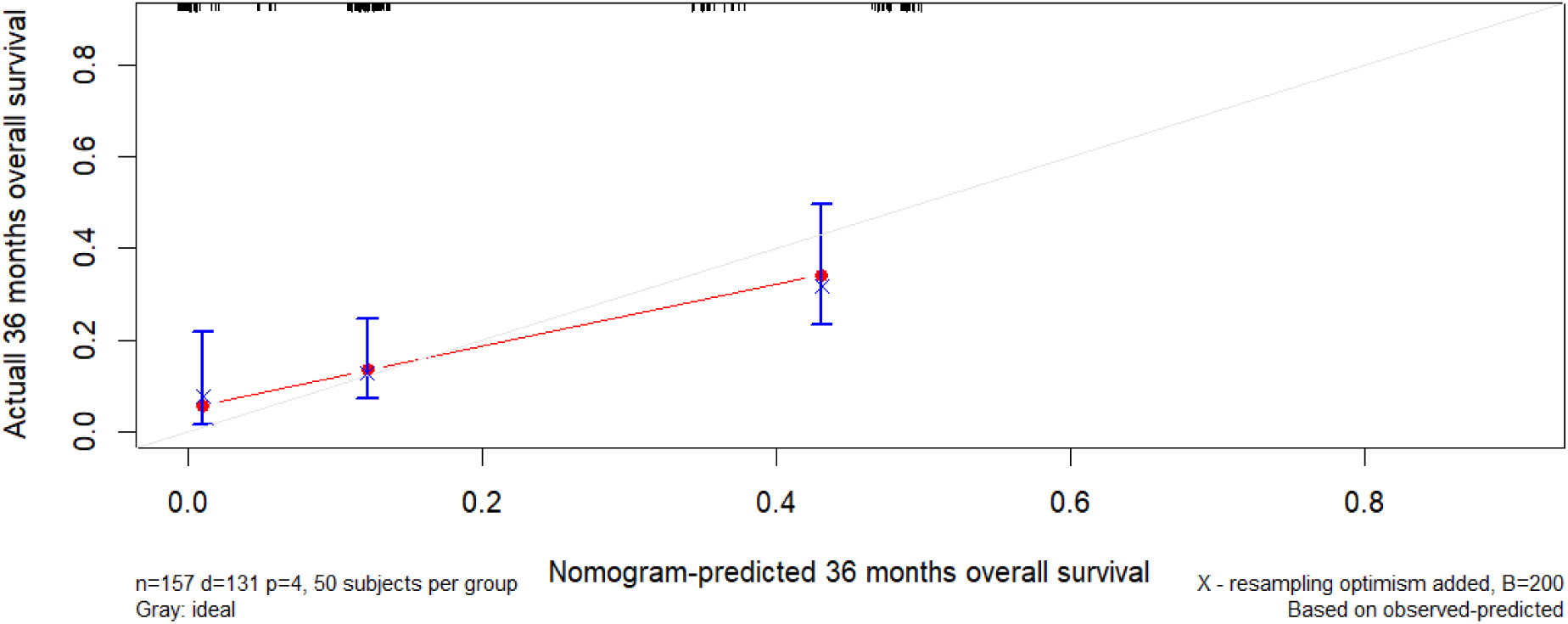
Calibration curve of the extended set 36 month nomogram

**Fig 34.**
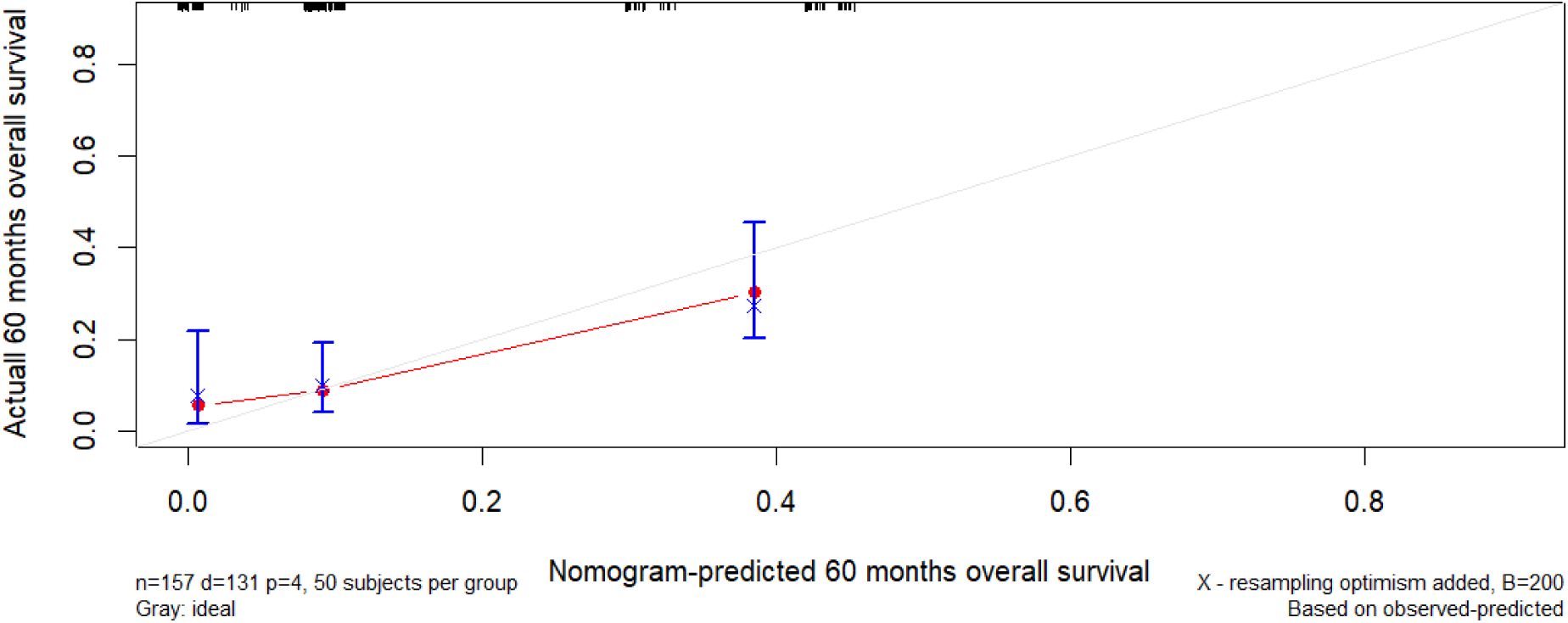
Calibration curve of the extended set 60 month nomogram

**Fig 35.**
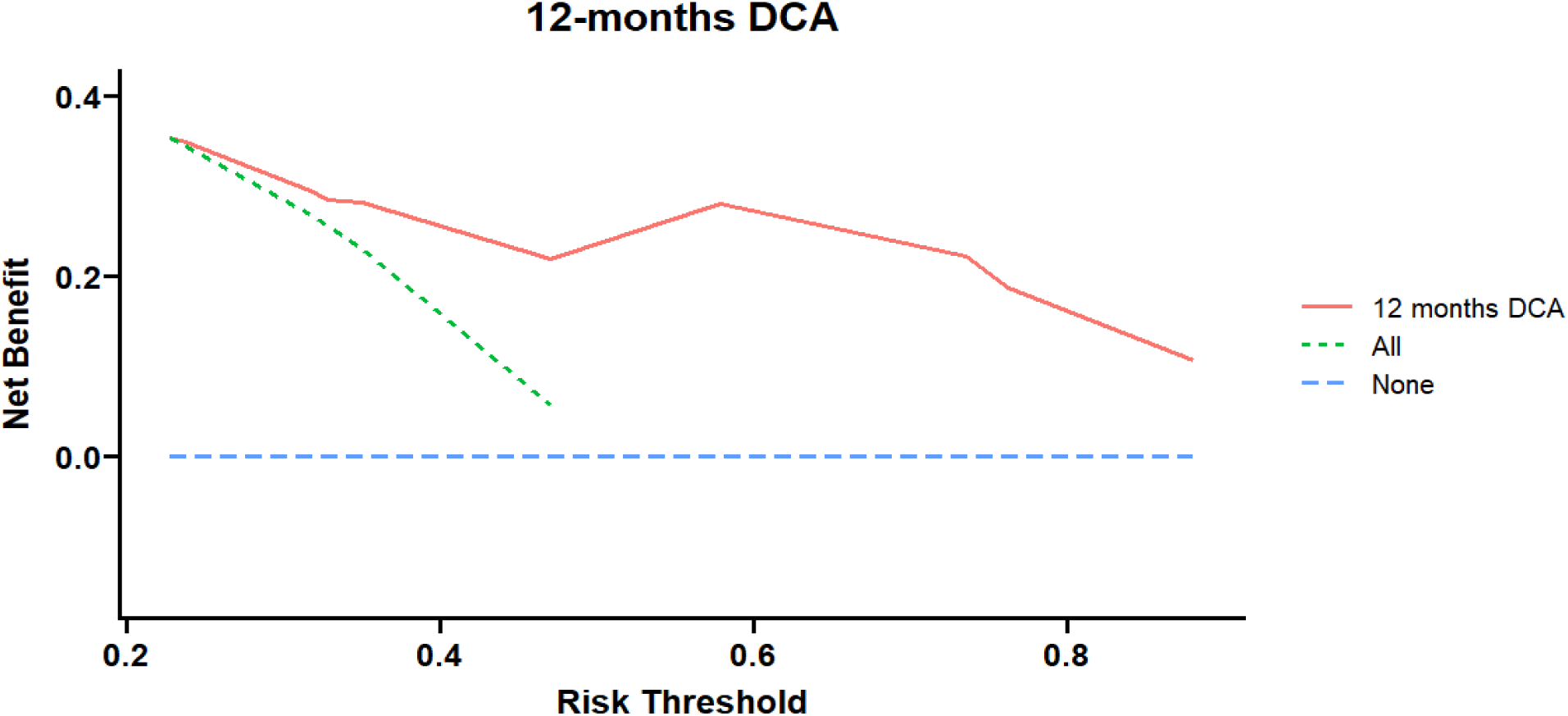
DCA curve for the extended set at 12 months

**Fig 36.**
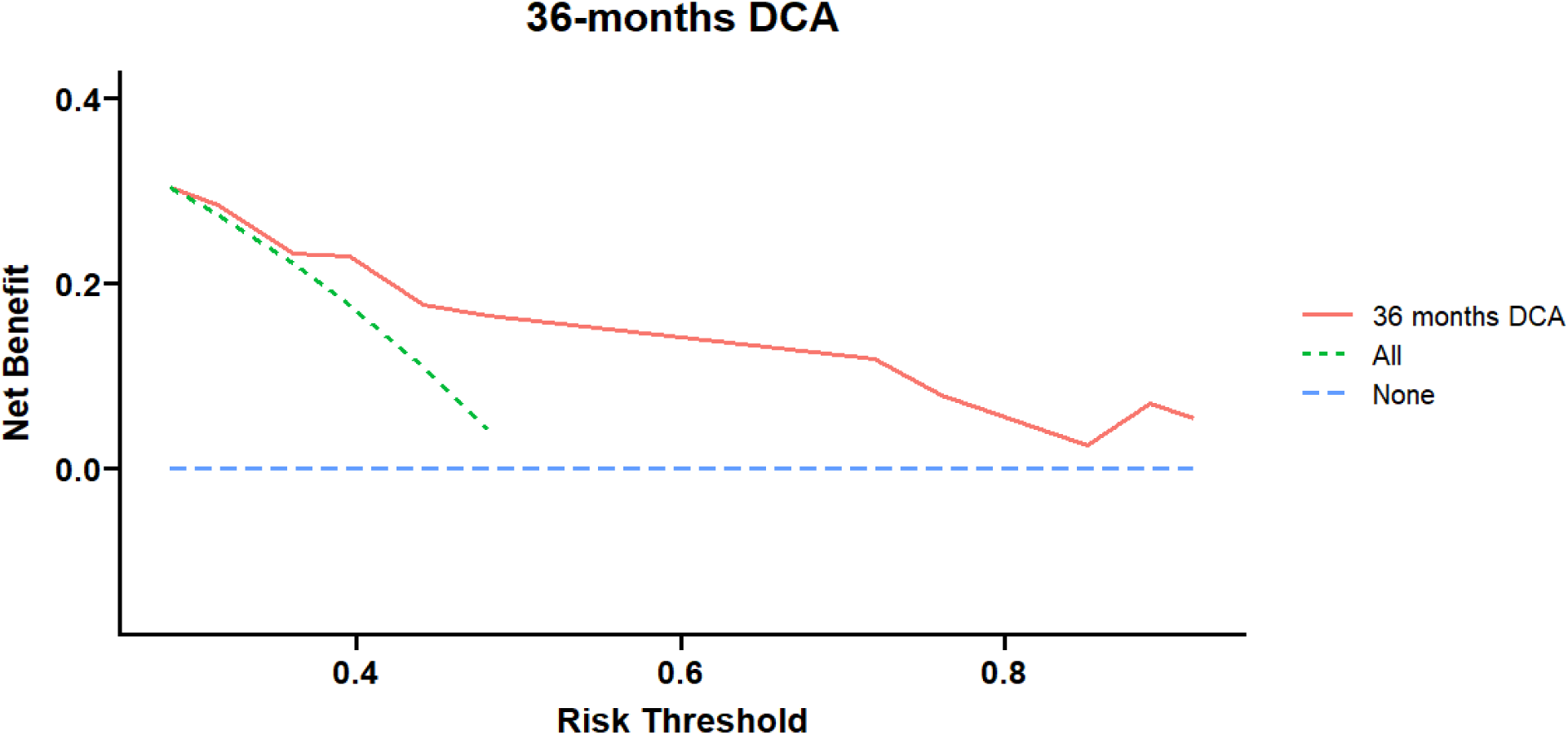
DCA curve for the extended set at 36 months

**Fig 37.**
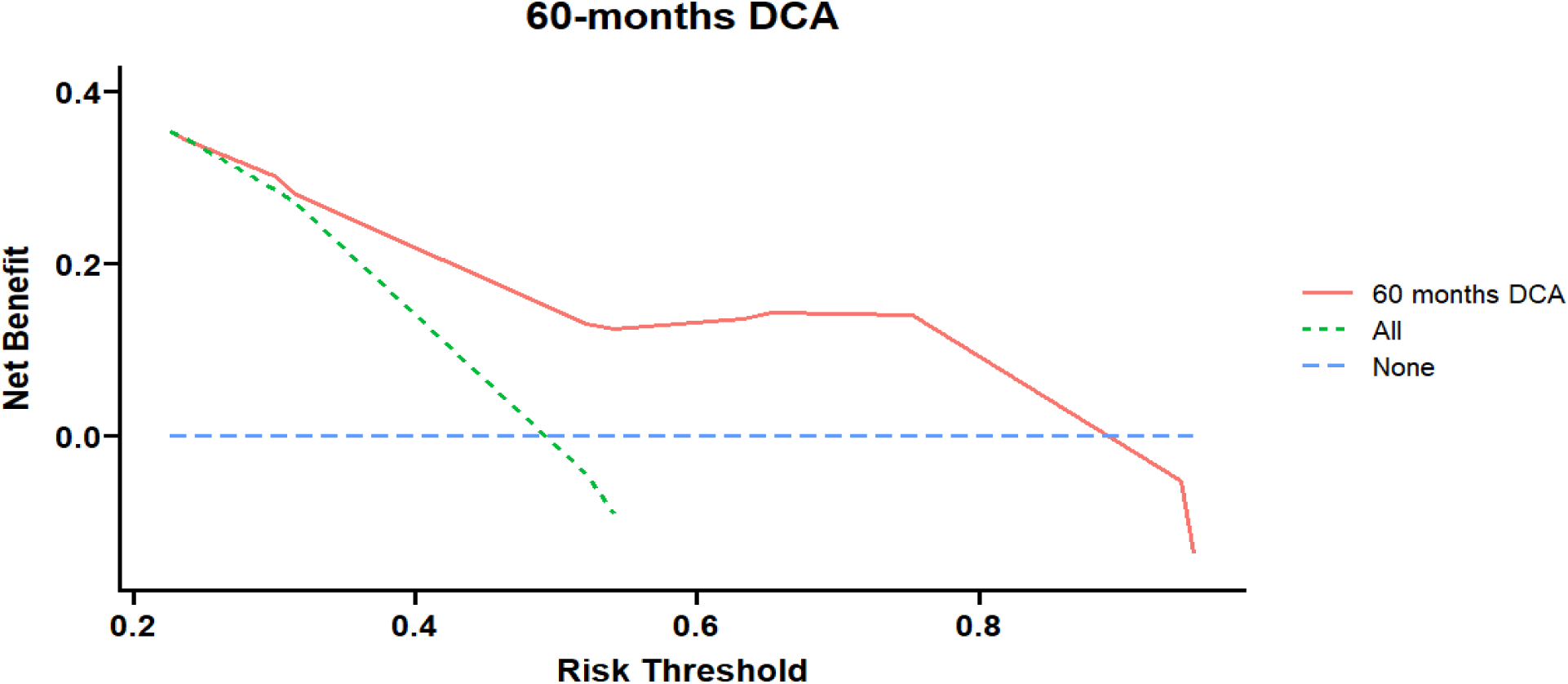
DCA curve for the extended set at 60 months

**Fig 38.**
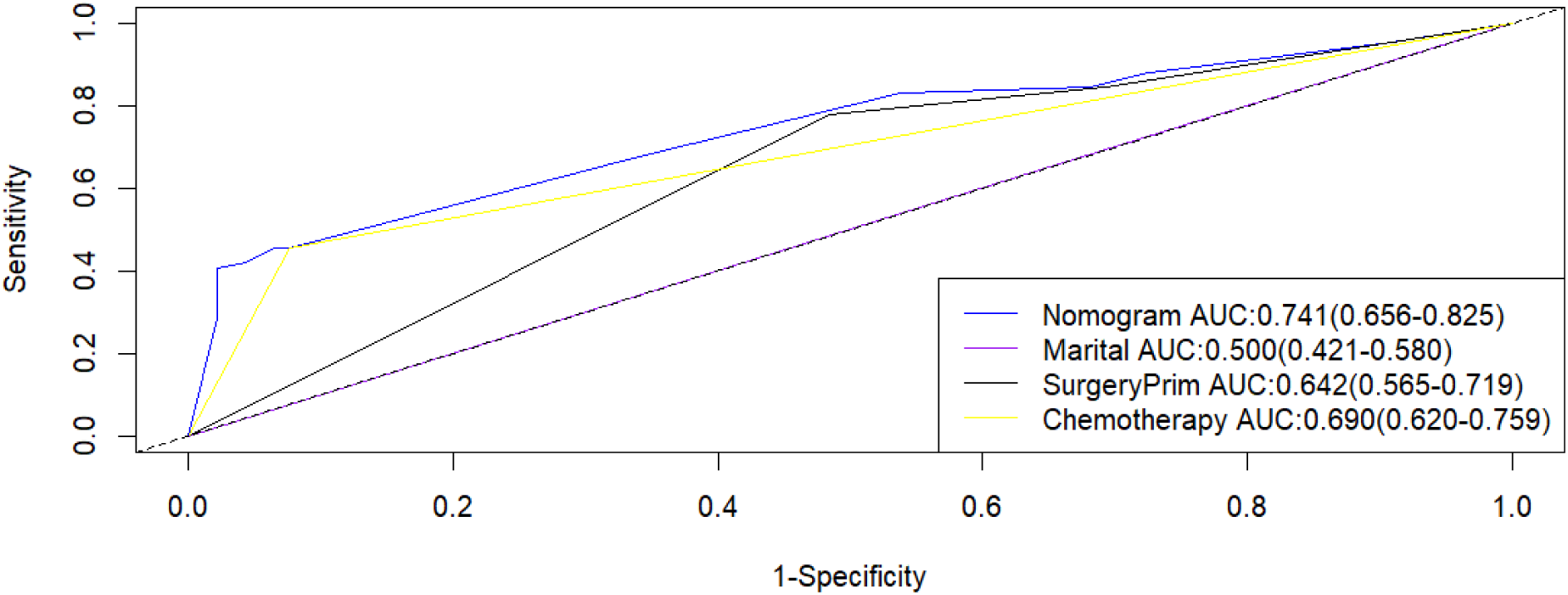
Comparison of the areas under the receiver operating characteristic curves for each factor at 12 months in the extended set

**Fig 39.**
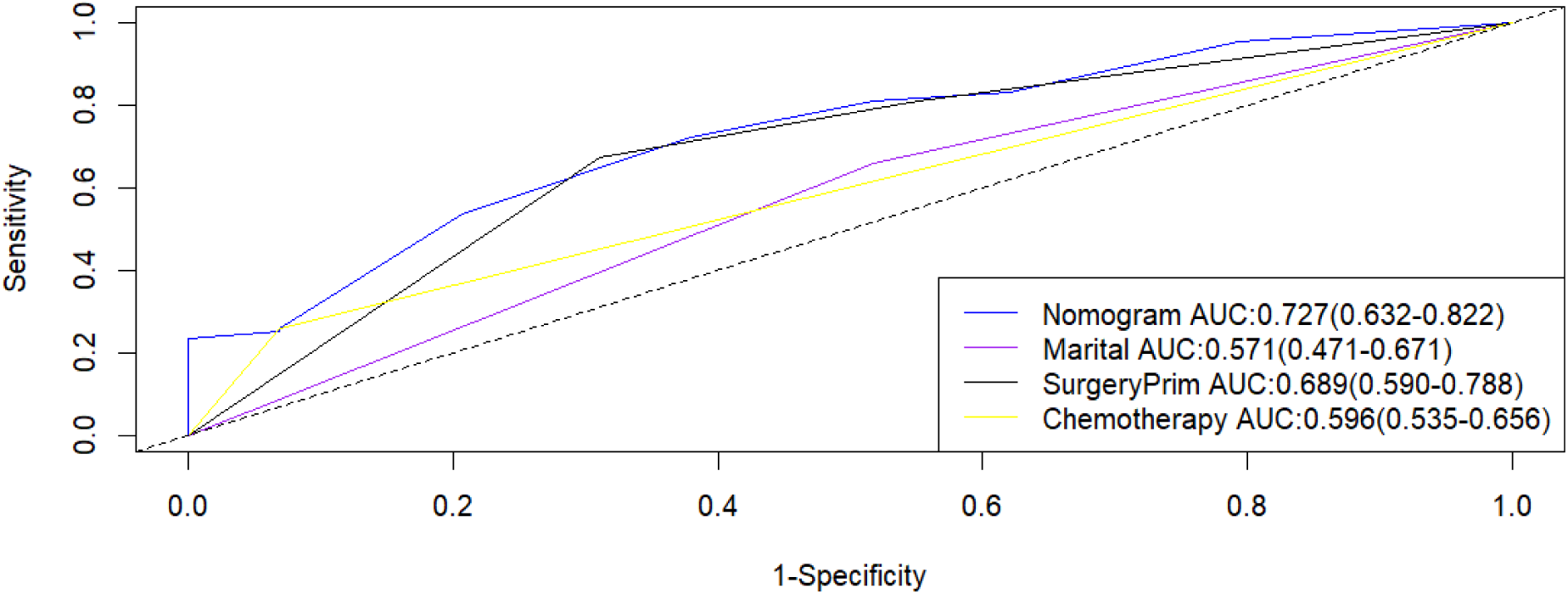
Comparison of the areas under the receiver operating characteristic curves for each factor at 36 months in the extended set

**Fig 40.**
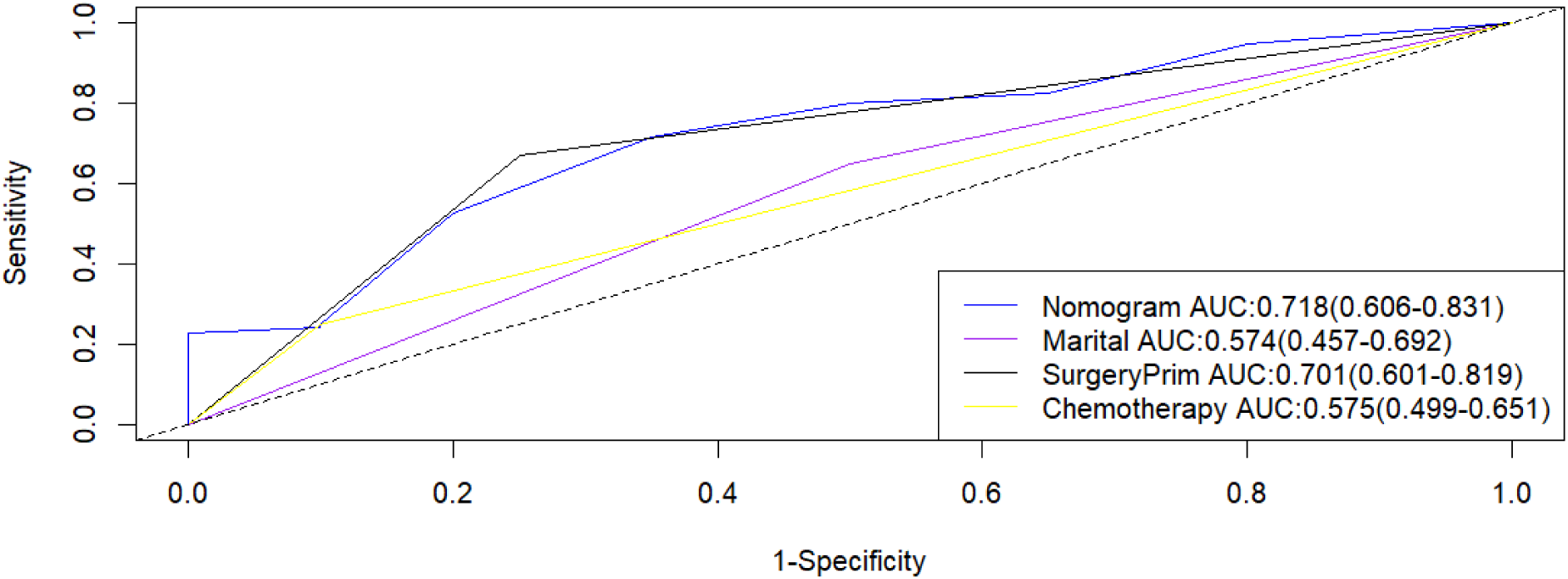
Comparison of the areas under the receiver operating characteristic curves for each factor at 60 months in the extended set

**Fig 41.**
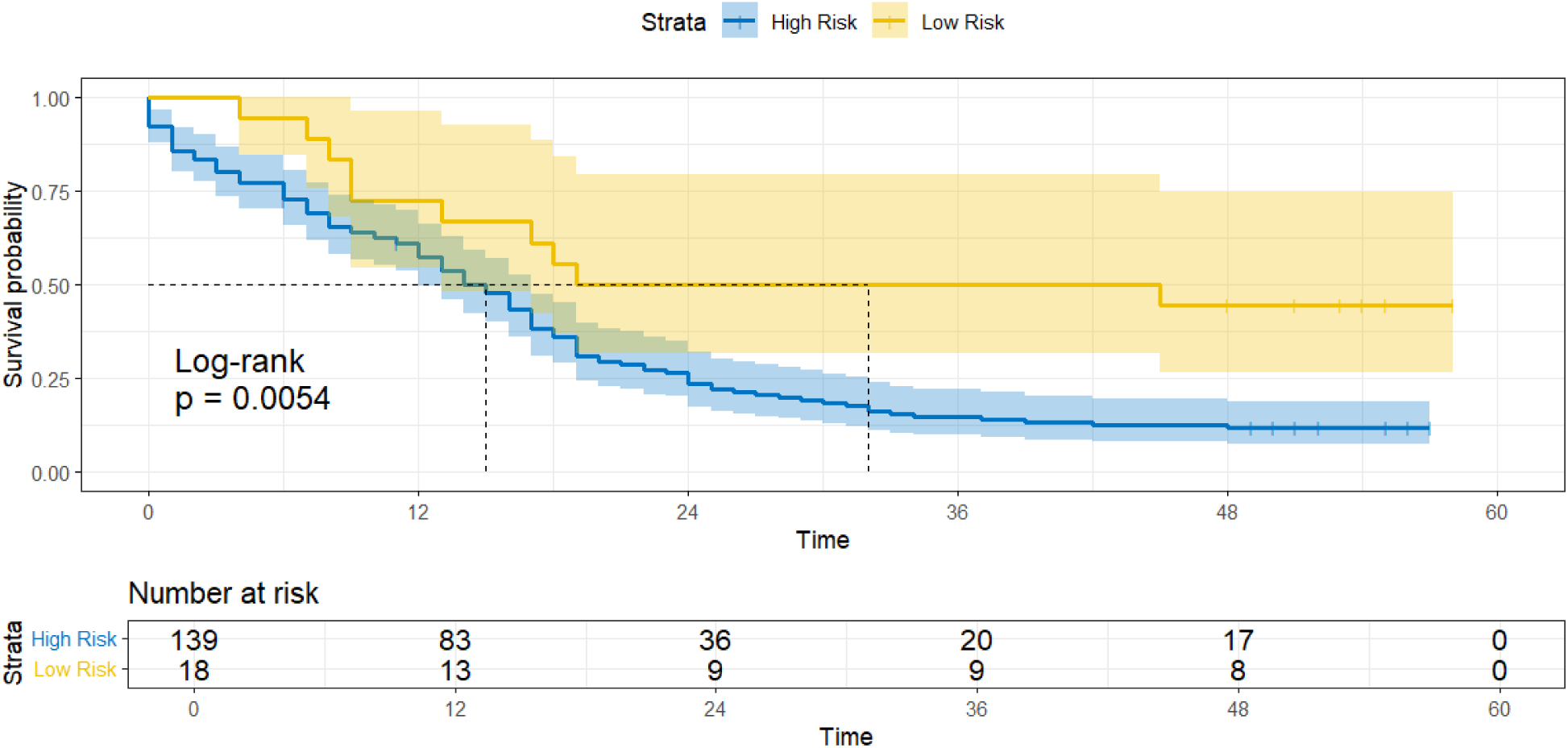
OS of high-risk and low-risk groups in the extended set

## Discussion

TNBC is a type of breast cancer with unique biological characteristics and clinical manifestations, which is prone to distant metastasis[13]. Due to the diversity and heterogeneity of TNBC, the prognosis and disease course should be adjusted according to the physiological and clinical characteristics of patients. In this study, we performed an in-depth analysis of risk factors for distant metastasis and prognosis after distant metastasis. Through the statistical analysis of the clinical data, we identified multiple factors that significantly affected the risk of distant metastasis and the prognosis. In this study, 16,959 eligible TNBC patients were screened using the SEER database. To further screen and analyze the data, we used univariate, multivariable logistic regression analysis in distant metastasis correlation factor analysis and Cox regression analysis in prognostic correlation factor analysis. This analysis showed that marital status, T stage, N stage, surgery, radiotherapy, and tumor size, were independent risk predictors of distant metastasis in patients with TNBC. Marital status, surgery, and chemotherapy are significant predictors of OS in patients with distant metastasis of TNBC. These independent risk variables were mostly consistent with clinical observations, and we constructed a diagnostic nomogram to predict distant metastasis in TNBC patients, and a prognostic nomogram in such patients. Relevant scores can be calculated by obtaining data from several key accessible variables on the nomogram, while our nomogram shows that we have a strong risk classification ability for TNBC patients that can be applied to patient survival information, as well as direct clinical decision-making and treatment allocation. We recommend classifying high-risk patients as high-risk patients according to tumor map, giving intensive care and thorough follow-up due to poor prognosis, and overall, using this model, investigators can reduce patient psychological distress, improve treatment and follow-up compliance, and collect more valuable data to design clinical trials.

In recent years, although similar work has been done by other researchers[14–16], most were performed at the molecular level rather than clinicopathological features. In this study, we combined the latest large sample and comprehensive clinical information from the SEER database and found that the incidence of distant metastasis was 4.3%, lower than the previous study, we identified seven important predictors of distant metastasis in triple-negative breast patients, namely, marriage, pathological type, N stage, T stage, radiotherapy, surgery and tumor size. Multiple prior reports have consistently demonstrated that younger age is closely linked to distant metastasis and unfavorable prognosis. A study encompassing 143 TNBC patients revealed that individuals aged 35 years or younger exhibited a 2.71-fold higher risk of relapse-free survival (RFS) and a 2.08-fold greater risk of distant relapse-free survival (DRFS) compared to those over 50 years of age [17]. Separately, another study enrolled 1930 TNBC patients, with 15% being diagnosed at an age below 40 and the remaining 85% being 40 years or older. Upon a median follow-up duration of 74 months, no significant disparities were observed in terms of local recurrence (LR) or disease-free survival (DFS) among different age groups. Multivariate analysis further highlighted that larger tumor size, lymphovascular invasion, and positive lymph node status were significant predictors of increased disease recurrence (DR) risk, whereas age and surgical approach did not exhibit any notable association with either outcome [18]. Furthermore, in another report with 254 patients with triple-negative breast cancer (TNBC), 75.6% of the patients were younger than 65 years old, and 24.4% were aged 65 or older. The results showed that the incidence of distant visceral metastases was significantly higher than that of bone metastases in both age groups (p<0.001). A significantly higher number of local recurrences, bone metastases, and secondary lymph node metastases were observed in the younger patients (p=0.035, 0.025, and 0.041, respectively) [19]. In contrast, our results suggest that age is not an independent risk factor for distant metastasis in patients with TNBC. The possible reasons for these differences are as follows. Our study specifically focused on TNBC rather not other breast cancer subtypes. We also recruited more TNBC patients, so the analytical power in our study seems more convincing.

This study identified marital status as an independent risk factor for distant metastasis and prognosis of primary TNBC. Some previous reports suggest that separated and widowed white women have a higher risk of death than married white women, and equally for single and divorced white women. For Asian / Pacific Islanders (API), only single and divorced women had a higher risk of death than married women. Marital status did not impact on the risk of death among Black or Hispanic women[20]. In the construction of another predictive model, patients with better survival outcomes were unmarried, uninsured, patients with higher T and N stages, and histological type of mixed invasive ductal carcinoma and lobular carcinoma. The prognosis of eTNBC patients is mainly influenced by marital status, insurance status, T stage, N stage and histological type, and the prognostic nomogram established based on these factors has high reliability[21]. In a nomogram used to predict the overall postoperative survival of Chinese TNBC patients, among 336 patients, age, marital status, tumor site, grade, T stage and N stage were independent prognostic factors[22]. Unmarried women are reported to be more likely to develop advanced breast cancer because having a spouse may be a factor in supporting early detection of malignancy[23,24]. In our study, it was found that being married was a relevant risk factor for developing a distant metastasis, while being unmarried was a related prognostic risk factor after having a distant metastasis.

Tissue type, T stage, N stage, and tumor size were shown to be relevant factors for distant metastasis in TNBC. Wang J et al[25] reported that in the nomogram of triple-negative breast cancer lung metastasis, age, tumor size, T stage and N stage were independent risk factors for LM in patients with TNBC. While the histological type, marital status, preoperative surgery, chemotherapy, bone metastasis, brain metastases, and LM were independent prognostic factors in patients with TNBC. In contrast to our study, histological type was associated with distant metastasis and not prognostic factors, which may be related to patient selection, and the literature has only selected patients with lung metastases.

Previous studies have shown that chemotherapy and surgery can significantly improve patient outcomes[26]. At present, chemotherapy remains the standard treatment for patients with TNBC[27]. Previous studies have shown that patients can benefit from surgery despite metastasis to distant organs[28,29]. Our prognostic nomogram indicates that surgery and chemotherapy are beneficial for the survival of patients with TNBC accompanied by distant metastasis. Therefore, identifying independent prognostic factors may help in identifying high-risk patients and establishing a specific surveillance program.

However, our study has several limitations. The SEER database contains data on the first diagnosis of the disease; therefore, metastases occurring in late stages cannot be recorded.

Furthermore, this is a retrospective study with a large sample size; therefore, selection bias is inevitable. The prognostic nomogram was constructed and validated at a single institution, which may partly influence its clinical applicability. Therefore, further calibration of the nomogram is needed in future studies.

## Conclusion

Our study identified that marital status, histological type, N stage, T stage, surgery, radiotherapy, and tumor size were independent risk factors for distant metastasis of TNBC, and that marital status, surgery, and chemotherapy were independent prognostic factors in patients with distant metastasis. We developed two nomograms utilizing diverse indicators to directly assess patient risk. The impressive performance of our nomograms in both the training and validation cohorts holds promise for assisting doctors in determining patient prognosis and devising personalized treatment plans.

